# Deciphering Colorectal Cancer-Hepatocyte Interactions: A Multiomic Platform for Interrogation of Metabolic Crosstalk in the Liver-Tumor Microenvironment

**DOI:** 10.1101/2024.12.06.627264

**Authors:** Alisa B. Nelson, Lyndsay E. Reese, Elizabeth Rono, Eric D. Queathem, Yinjie Qiu, Braedan M. McCluskey, Alexandra Crampton, Eric Conniff, Katherine Cummins, Ella Boytim, Senali Dansou, Justin Hwang, Sandra Safo, Patrycja Puchalska, David K. Wood, Kathryn L. Schwertfeger, Peter A. Crawford

## Abstract

Metabolic reprogramming is a hallmark of cancer, enabling tumor cells to adapt to and exploit their microenvironment for sustained growth. The liver is a common site of metastasis, but the interactions between tumor cells and hepatocytes remain poorly understood. In the context of liver metastasis, these interactions play a crucial role in promoting tumor survival and progression. This study leverages multiomics coverage of the microenvironment via liquid chromatography and high-resolution, high-mass accuracy mass spectrometry-based untargeted metabolomics, ^13^C-stable isotope tracing, and RNA sequencing to uncover the metabolic impact of co-localized primary hepatocytes and a colon adenocarcinoma cell line, SW480, using a 2D co-culture model. Metabolic profiling revealed disrupted Warburg metabolism with an 80% decrease in glucose consumption and 94% decrease in lactate production by hepatocyte-SW480 co-cultures relative to SW480 control cultures. Decreased glucose consumption was coupled with alterations in glutamine and ketone body metabolism, suggesting a possible fuel switch upon co-culturing. Further, integrated multiomic analysis indicates that disruptions in metabolic pathways, including nucleoside biosynthesis, amino acids, and TCA cycle, correlate with altered SW480 transcriptional profiles and highlight the importance of redox homeostasis in tumor adaptation. Finally, these findings were replicated in 3-dimensional microtissue organoids. Taken together, these studies support a bioinformatic approach to study metabolic crosstalk and discovery of potential therapeutic targets in preclinical models of the tumor microenvironment.

## Introduction

The liver is the site of one of six cancers with increasing incidence of primary tumors – hepatocellular carcinoma (hepatoma) (1). Moreover, the liver is a common site of solid tumor cell metastasis, including from breast and colorectal cancers, causing significant morbidity and mortality (2). Tumor cells within the liver interact with liver resident cell types, including hepatocytes. Metabolic adaptation is essential for tumor success throughout cancer cell transformation, proliferation, and metastasis (3, 4). Cancer cells reprogram their metabolism to meet increased demands for energy, biosynthesis, and redox homeostasis. Studies show this adaptation extends to the tumor microenvironment where cancer cells can tune and exploit their environment to meet metabolic needs (5–8). Within the liver, hepatocytes engage dynamic metabolic programs to support local and systemic physiology. Exploitation of this rich metabolic environment may represent an essential interaction that facilitates tumor cell colonization in the liver niche. However, our understanding of the specific interactions between cancer cells and hepatocytes that drive survival and proliferation in the metastatic liver niche remains limited. Previous studies profiling metabolism of HCC indicates transformed hepatocytes engage in aerobic glycolysis and altered lipid and amino acid metabolism (9, 10). Moreover, substrate fuels are used to program neighboring immune cells for repressed anti-tumor responses (7). Metastasizing cells have been shown to require adaptations that help them overcome the hypoxic liver microenvironment (11–13). But these studies do not consider the role of co-localized, non-transformed hepatocytes. Additionally, these studies are limited in their coverage of the omics landscape.

Metabolomics technologies are well-positioned to reveal potential metabolic adaptation in cancer. Mass spectrometry-based untargeted metabolomics surveys global metabolic shifts among samples by measuring the fluctuations of multiple chemical feature abundances detected as mass-to-charge (*m/z*) signal and retention time pairs (14–16). An additional dimension of information can be gained by the convergent use of ^13^C-labeled stable isotope tracers. While static measurement of metabolites provides a snapshot of perturbed influxes or effluxes that lead toward or away from measured metabolites, these measurements often fail to reveal nodes through which shifts occur without significant fold changes in the static abundance of metabolites (17). Merging the advantages of high-resolution mass spectrometry-based untargeted metabolomics with ^13^C-stable isotope labeling known as isotope tracing untargeted metabolomics (ITUM) provides a unique opportunity to discover dynamic and potentially crucial metabolic pathways (18–21). However, studying metabolic communication in the microenvironment is an ongoing challenge (22). Stable isotopes present an advantage, but developing model systems amenable to ITUM approaches while maintaining physiological relevance is difficult. Here, we present an approach using our untargeted metabolomics and ITUM pipelines in an engineered *in vitro* model to discover metabolic interactions between colon adenocarcinoma cells (SW480 cell line) and primary hepatocytes. To adapt our pipeline to a mixed cell model, we quantify extracellular and intracellular metabolite pools upon co-culture to identify impacted metabolic pathways. Additionally, we hypothesized that the combined study of differential glucose utilization and metabolite-metabolite relationships could reveal tumor cell metabolic adaptation in the hepatocyte-SW480 microenvironment. Therefore, we present approaches that extend the application of untargeted metabolomics and ITUM to reveal nodes of adaptation in the tumor-hepatocyte microenvironment. Finally, a multiomic integration with the transcriptome links metabolic adaptations to altered functional programs in tumor cells with co-localized hepatocytes.

## Results

### Co-culture of SW480 cells with primary hepatocytes reprograms metabolism

We used a co-culture model to study metabolic adaptation of tumor cells to the hepatocyte microenvironment. Culturing primary hepatocytes is challenged by their loss of hepatocyte-like function through de-differentiation (23, 24). However, co-culture with 3T3-J2 murine embryonic fibroblasts can sustain hepatocellular function for more than 6 weeks (25). Therefore, we directly co-cultured primary rat hepatocytes, murine 3T3-J2 fibroblasts (J2s), and the human colon adenocarcinoma cell line, SW480 (**Figure 1A**). To form 2D co-cultures, J2s were growth arrested and then plated with hepatocytes in 12-well plates to sustain hepatocellular function (HJ cultures). Control plates of J2s were plated on the same day in preparation for SW480-J2 (SJ) control co-cultures. After 7 days, SW480s were seeded to form SW480-J2-Hepatocyte (SJH) co-cultures and SJ controls. All media and cells were collected on day 10 for metabolomics and transcriptomics analyses (**Figure 1B**). In each experiment, the group of interest, SJH, was compared to HJ and SJ controls.

**Figure 1.**
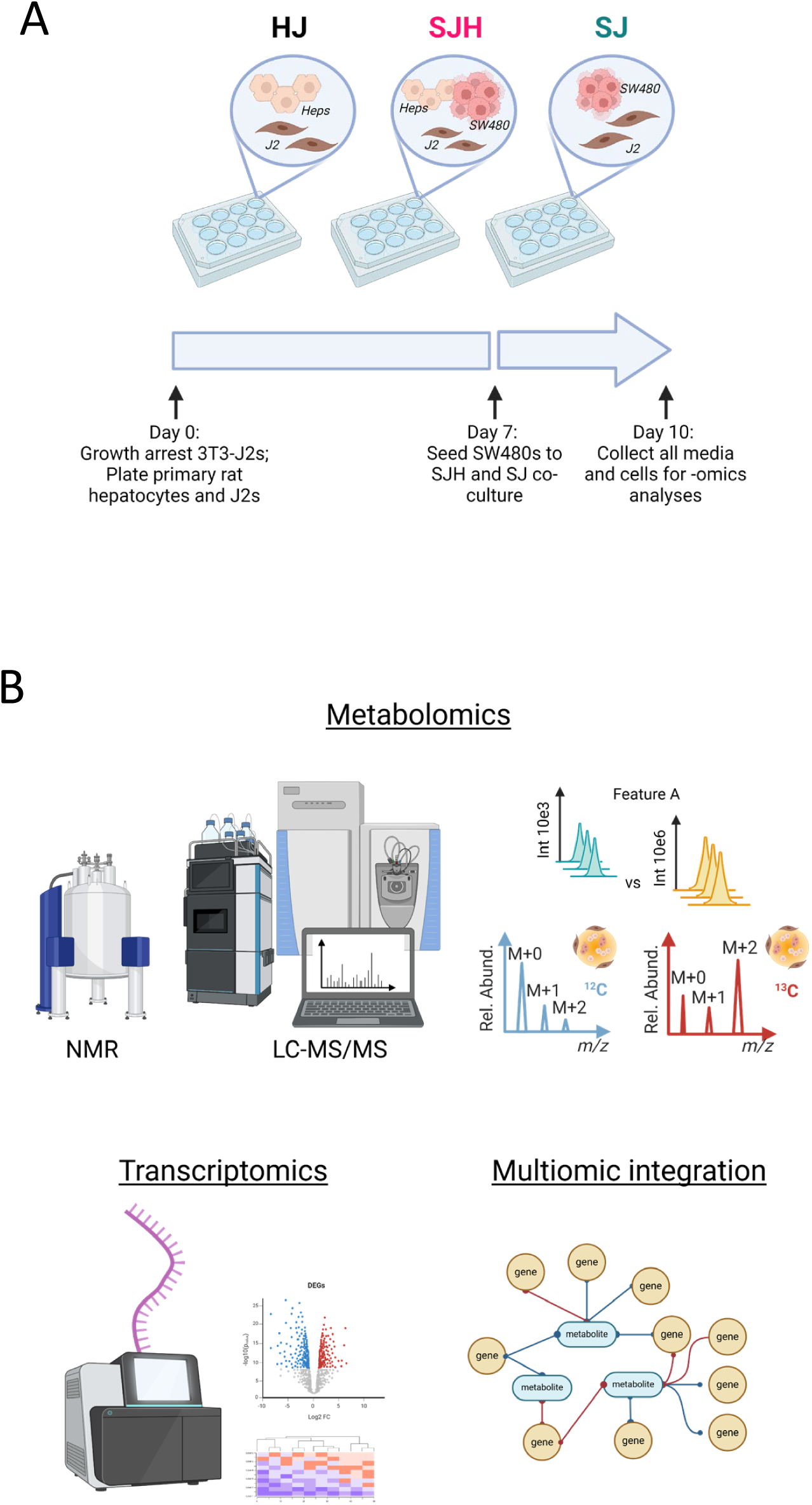
Multiomic study of co-cultures of primary hepatocytes and SW480s. (A) Scheme of 2D co-culture system and timeline for cell collection. (B) -Omic coverage of co-cultured groups.

Cancer metabolism is hallmarked by high glucose consumption and utilization through aerobic glycolysis, resulting in high lactate production, known as the Warburg Effect (26, 27). Hepatocytes play significant roles in regulating glucose homeostasis. We hypothesized that hepatocytes may impact glucose metabolism in co-cultures. Therefore, we quantified net changes in glucose and lactate concentrations in conditioned cell culture media using ^1^H- nuclear magnetic resonance (NMR) spectroscopy. Then, we normalized changes in exogenous metabolite concentrations to cellular biomass (DNA content) to report net glucose consumption and lactate production over 24 hours. As expected, SJ controls model the Warburg effect in culture, consuming 2.50 ± 0.01 mol glucose/day/mg DNA, and producing 3.86 ± 0.05 mol lactate/day/mg DNA (**Figure 2A**). The presence of hepatocytes decreased total glucose consumption by 80% (0.5 ± 0.05 mol glucose/day/mg DNA) and lactate production by 94% to 0.23 ± 0.01 mol lactate/day/mg DNA. Furthermore, the ratio of lactate to glucose significantly decreased from 1.54 ± 0.02 to 0.46 ± 0.05, suggesting the presence of hepatocytes disrupts Warburg metabolism of SW480s (**Figure 2B**).

**Figure 2.**
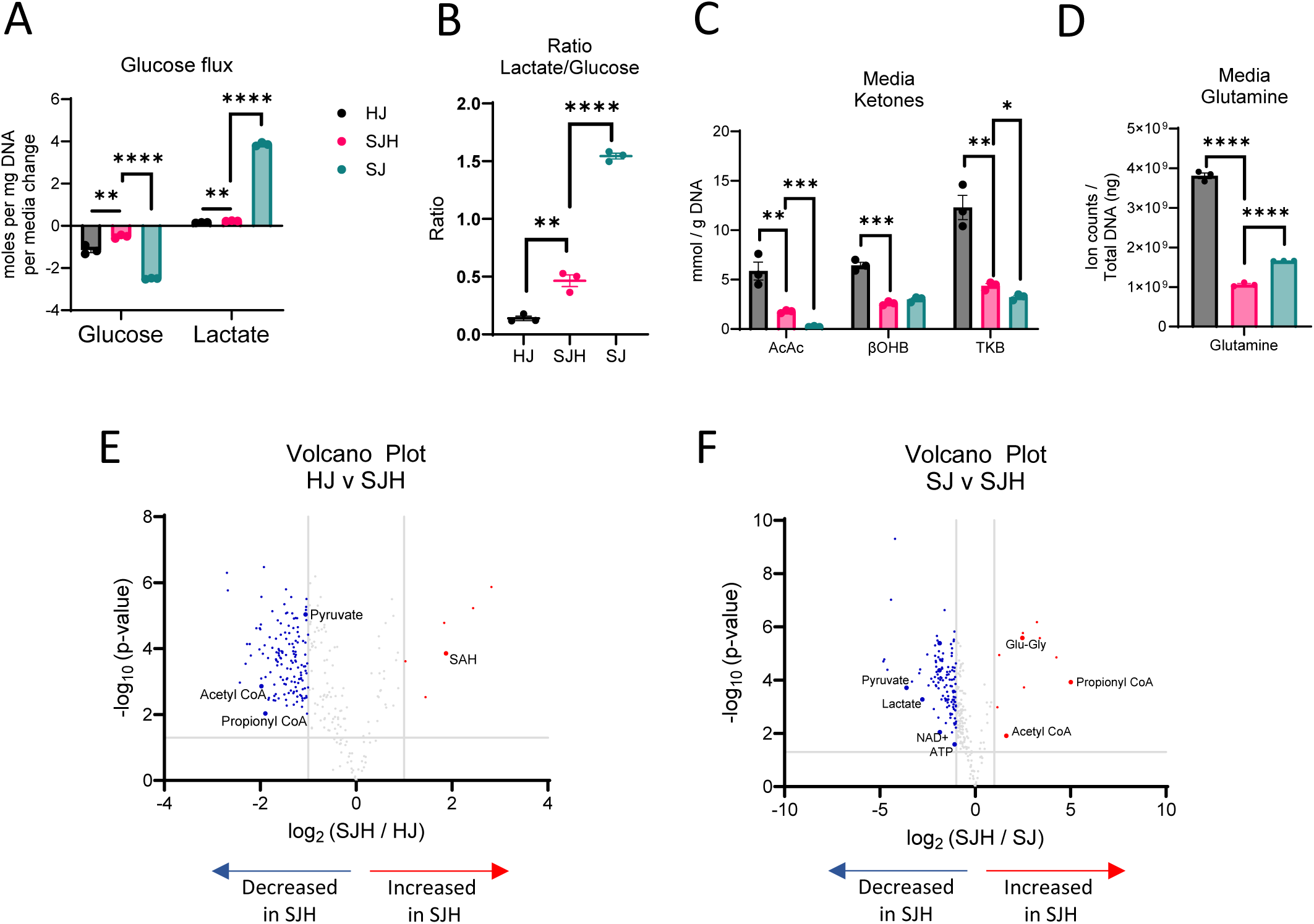
Fuel utilization in 2-dimensional co-cultures. (A) Glucose consumption and lactate production in moles per ng total DNA per day. (B) Lactate / glucose ratio of media concentration after 24h. (C) Concentration of acetoacetate (AcAc), β-hydroxybutryrate (βOHB) and total ketone bodies (TKB) in mmol/g DNA measured in media after 24h incubation. (D) Relative abundance of glutamine in total ion counts after normalization to total ng DNA. Volcano plot showing upregulated and downregulated metabolites in SJH compared to: (E) HJ control cultures and (F) SJ control cultures; positive log_2_FC = up in SJH. Significance tested using unpaired t-test, comparison HJ vs SJH or SJ vs SJH, and corrected for multiple comparisons using Benjamini-Hochberg method. *: p adj. < 0.05, **: p adj. < 0.01, ***: p adj. < 0.001, ****: p adj. < 0.0001. Abbreviations: HJ: Hepatocyte+3T3-J2 co-culture; SJH: SW480+3T3-J2+Hepatocyte co-culture; SJ: SW480+3T3-J2 co-culture.

To determine if decreased utilization of glucose by SW480 cells when co-cultured with hepatocytes could result from a fuel switch from glucose to other available substrates, we quantified hepatocyte-derived ketones and culture media-derived glutamine. Using a UHPLC-MS/MS based approach, we quantified total ketone bodies in the conditioned media of co-cultures after 24 hours. As expected, primary rat hepatocyte control co-cultures produced the ketone bodies acetoacetate (AcAc) and β-hydroxybutyrate (βOHB, **Figure 2C**). In the presence of SW480s, total ketone bodies recovered in the media were diminished by 64%, suggesting either decreased production by hepatocytes or increased consumption of ketones by non-hepatocyte cells. Glutamine abundance was also significantly decreased in SJH co-cultures relative to both HJ and SJ controls (**Figure 2D**). Interestingly, glutamine abundance decreased 36% in SJH relative to SJ controls, suggesting the presence of hepatocytes further enhanced glutamine dependence of SW480s. Together these data suggest an adaptation to fuel utilization in hepatocyte-SW480 co-cultures.

To survey metabolic interaction-dependent metabolite shifts resulting from co-culture, we next used differential analysis of features detected by LC-MS-based label-free untargeted metabolomics. Prior to any downstream analysis, low variance features were removed, features of interest were annotated in Compound Discoverer 3.3, and when possible, matched to commercial standards and MS/MS spectral libraries. Five metabolic features by LC-MS-based untargeted profiling increased more than 2-fold (log_2_ ≥ 1) in SJH compared to HJ controls. This included putative S-adenosylhomocysteine (SAH; **Table 1**, **Figure 2E**), an intermediate of one-carbon metabolism. Compared to HJ, SJH co-cultures showed 142 downregulated features, including pyruvate (**Supplemental Figure 1A**), acetyl-CoA (**Supplemental Figure 1B**), and propionyl-CoA (**Supplemental Figure 1C**), metabolites important in glucose metabolism and the TCA cycle. Relative to SJ controls, SJH co-cultures show a greater than 2-fold increase in 10 features, including propionyl-CoA (**Figure 2F; Supplemental Figure 1C**), glutamyl-glycine, an intermediate of glutathione metabolism, and acetyl-CoA (**Supplemental Figure 1B**). A ≥2-fold decrease was observed in 119 features in SJH compared to SJ controls, including pyruvate (**Supplemental Figure 1A**), lactate (**Supplemental Figure 1D**), NAD^+^ (**Supplemental Figure 1E**), and ATP (**Supplemental Figure 1F**). However, decreases in the NAD^+^ pool did not lead to a significant change in the NAD^+^/NADH ratio in SJH relative to SJ (**Supplemental Figure 1G**). Finally, we calculated energy charge to determine the current energy status of the co-culture based on relative abundance of AMP, ADP, and ATP pools. We saw a modest, but not statistically significant, decrease in SJH relative to SJ co-cultures, suggesting a possible decrease in available energy (**Supplemental Figure 1H-I**).

Untargeted metabolomics of mixed cell populations represent metabolite pools that are combined across all cell types. Therefore, changes in relative abundance may represent an adaptation in one cell type, a combination of adaptations across multiple cell types, or could simply be a product of dilution of the pool after adding biomass. Due to the observed number of features that were significantly decreased in SJH relative to either HJ or SJ controls, we sought to determine if our approach detects true biological interactions in hepatocyte and SW480 co-cultures, rather than simple dilutions of HJ or SJ metabolite pools. Therefore, we implemented a dilution approach to identify those features that were significantly different from a HJ-SJ dilution (**Figure 3A**). HJ and SJ extracts were combined 1:1 (1T1) prior to LC-MS injection, alongside HJ, SJ, and SJH samples. Putative metabolites with pool sizes that significantly differed between SJH and controls were compared to the 1T1 dilution sample. A low threshold for discovery was used (p< 0.05) to identify those features that may indicate metabolic interactions. Lactate was significantly decreased in SJH relative to 1T1 samples (**Figure 3B**; **Table 2**). Normalized ion counts for each group and the analytical control, 1T1, are shown in **Supplemental Figure 2**, demonstrating a likely metabolic interaction between SW480s and hepatocytes beyond an outcome that could be explained by simple metabolite pool dilution. Additionally, four glutamyl peptides – including glutamyl-glycine, as well as uracil, uridine diphosphate (UDP), aspartate, and malate were increased more than 2-fold in SJH relative to 1T1 (**Figure 3B**; **Table 2**). Employment of an analytical 1T1 dilution of controls for differential analysis of directly co-cultured cells reveals biological derangement of metabolite abundances belonging to glycolytic, amino acid, and nucleoside pathways.

**Figure 3.**
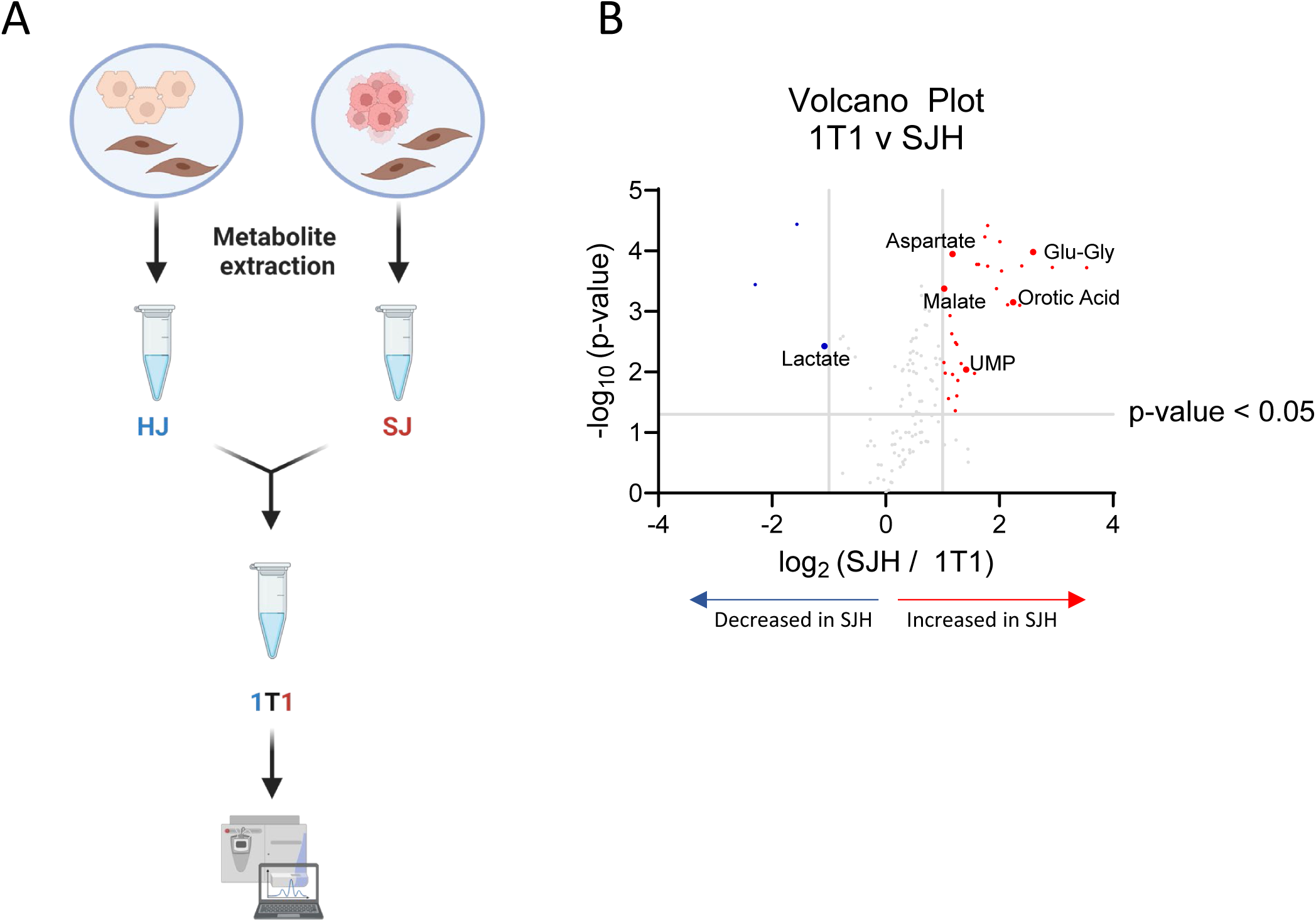
Analytical dilution of co-culture controls reveals metabolic adaptation in SJH co-cultures. (A) Schematic of analytical dilution of HJ and SJ controls to form a 1-to-1 ratio (1T1) after metabolite extraction. (B) Volcano plot upregulated and downregulated metabolites in SJH compared to 1T1 ion counts; positive log_2_FC = up in SJH.

**Table 1.** Differential abundance of select metabolites in SJH co-cultures relative to HJ or SJ controls. FC: fold change; ctrl: control.

| Putative metabolite | Log<sub>2</sub> (FC: SJH / ctrl) | Adjusted p-value | Direction;<br>Relative to [HJ] or [SJ] |
| --- | --- | --- | --- |
| S-Adenosylhomocysteine | 1.87 | 0.0004 | ↑; HJ |
| Pyruvate | -1.05 | 0.0001 | ↓; HJ |
| Acetyl-CoA | -1.98 | 0.002 | ↓; HJ |
| Propionyl-CoA | -1.89 | 0.012 | ↓; HJ |
| Propionyl-CoA | 5.01 | 0.0004 | ↑; SJ |
| Glutamyl-glycine | 2.47 | 0.00007 | ↑; SJ |
| Acetyl-CoA | 1.63 | 0.016 | ↑; SJ |
| Pyruvate | -3.60 | 0.0006 | ↓; SJ |
| Lactate | -2.78 | 0.0012 | ↓; SJ |
| NAD<sup>+</sup> | -1.85 | 0.012 | ↓; SJ |
| ATP | -1.09 | 0.03 | ↓; SJ |

**Table 2.** Differential abundance of select metabolites in SJH co-cultures relative to 1T1 dilution. Abbreviations: FC: fold change.

| Putative metabolite | Log<sub>2</sub> (FC: SJH / HJ) | Adjusted p-value | Direction relative to 1T1 |
| --- | --- | --- | --- |
| Glutamyl-glycine | 2.59 | 0.0018 | ↑ |
| Uracil | 2.24 | 0.0044 | ↑ |
| Uridine diphosphate | 1.42 | 0.021 | ↑ |
| Aspartate | 1.17 | 0.0018 | ↑ |
| Malate | 1.03 | 0.0029 | ↑ |
| Lactate | -1.08 | 0.013 | ↓ |

### Metabolite exchange of nucleoside intermediates

Untargeted metabolomics results indicate altered activity in both purine and pyrimidine pathways when SW480s are co-cultured with hepatocytes, whose identities were confirmed by retention time and MS/MS fragmentation match to commercial standards (**Figure 2E-F**; **Figure 3B**). We hypothesized that the presence of hepatocytes may facilitate adaptation through exchange of purine and pyrimidine intermediates. Therefore, we performed LC-MS-based untargeted metabolomics on cell extracts and media after the first 24 hours of direct co-culture to evaluate fold changes in nucleoside intermediate abundance over time. In all three groups, hypoxanthine, an intermediate of purine metabolism, was depleted comparing starting media (t0) to media harvested after 24 hours, indicating uptake from serum-containing media or conversion to other metabolic products (**Figure 4A**, **Table 3**). SJH co-cultures had 23% (±3%) greater depletion of hypoxanthine compared to HJ controls and 12% (±4%) less than SJ controls (**Supplemental Figure 3**). Interestingly, hypoxanthine abundance was coupled with diminished extracellular uric acid, the terminal product of purine degradation, in HJ and SJH groups, while SJ controls show an average 809% increase in uric acid (**Figure 4A, Supplemental Figure 3**). Hypoxanthine and uric acid are both metabolites that can be found in starting media due to the presence of serum. We further analyzed media under normal culture conditions and in the absence of cells to control for possible spontaneous degradation and observed an accumulation of hypoxanthine and uric acid (9% and 47%, respectively) after 24 hours, suggesting instability of upstream metabolites at 37°C (denoted as dotted lines, **Supplemental Figure 3**). Therefore, these data indicate significant metabolic activity in the purine degradation pathway in SJ controls that is altered by the presence of hepatocytes. To investigate if this impacts intracellular purine intermediates, we measured inosine in cellular extracts. Inosine, an intermediate in the purine salvage pathway, showed a 2-fold increase in hepatocyte-containing cultures while it was modestly diminished in SJ controls after 24 hours (**Figure 4B**). Finally, inosine was uniquely enriched from [U-^13^C_6_]glucose in HJ and SJH groups but not detectable in SJ in 3D microtissue organoids (**Figure 4C**). Conversely, the average total ^13^C-enrichment of HJ and SJH groups was 58.7% (± 1.5%) and 64.7% (± 4.5%) of the total inosine pool. This observation suggests that changes in glucose contribution to inosine pools may be localized to hepatocytes. Together these data suggest the presence of hepatocytes increases purine salvage, rescuing cells from uric acid accumulation.

**Figure 4.**
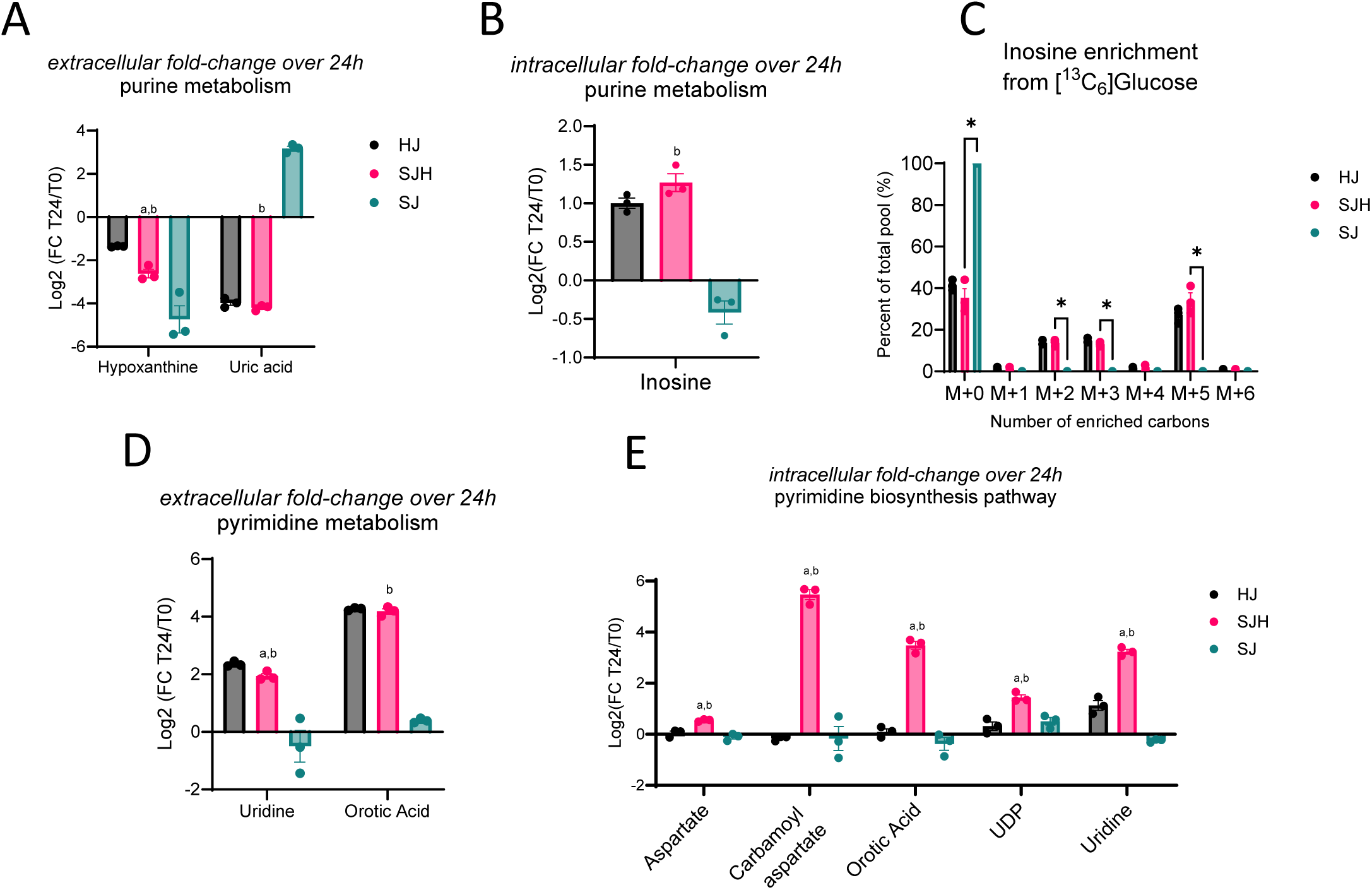
Metabolic interactions of biosynthetic pathways in SJH co-cultures. Fold change of metabolite abundance after 24h co-culture relative to time point 0 in: (A) media abundance of purine metabolism products, hypoxanthine and uric acid, (B) intracellular inosine pools, (D) media abundance of pyrimidine biosynthesis intermediates, uridine and orotic acid, and (E) intracellular pyrimidine intermediates and substrates, aspartate, carbamoyl aspartate, orotic acid, UDP, and uridine. (C) ^13^C-enrichment of intracellular inosine pools from 22 mM [U-^13^C_6_]glucose in 3D microtissue organoids. Statistical comparison by unpaired t-test; letters indicate significance in comparison to HJ controls (“a”) or SJ controls (“b”).

**Table 3.**
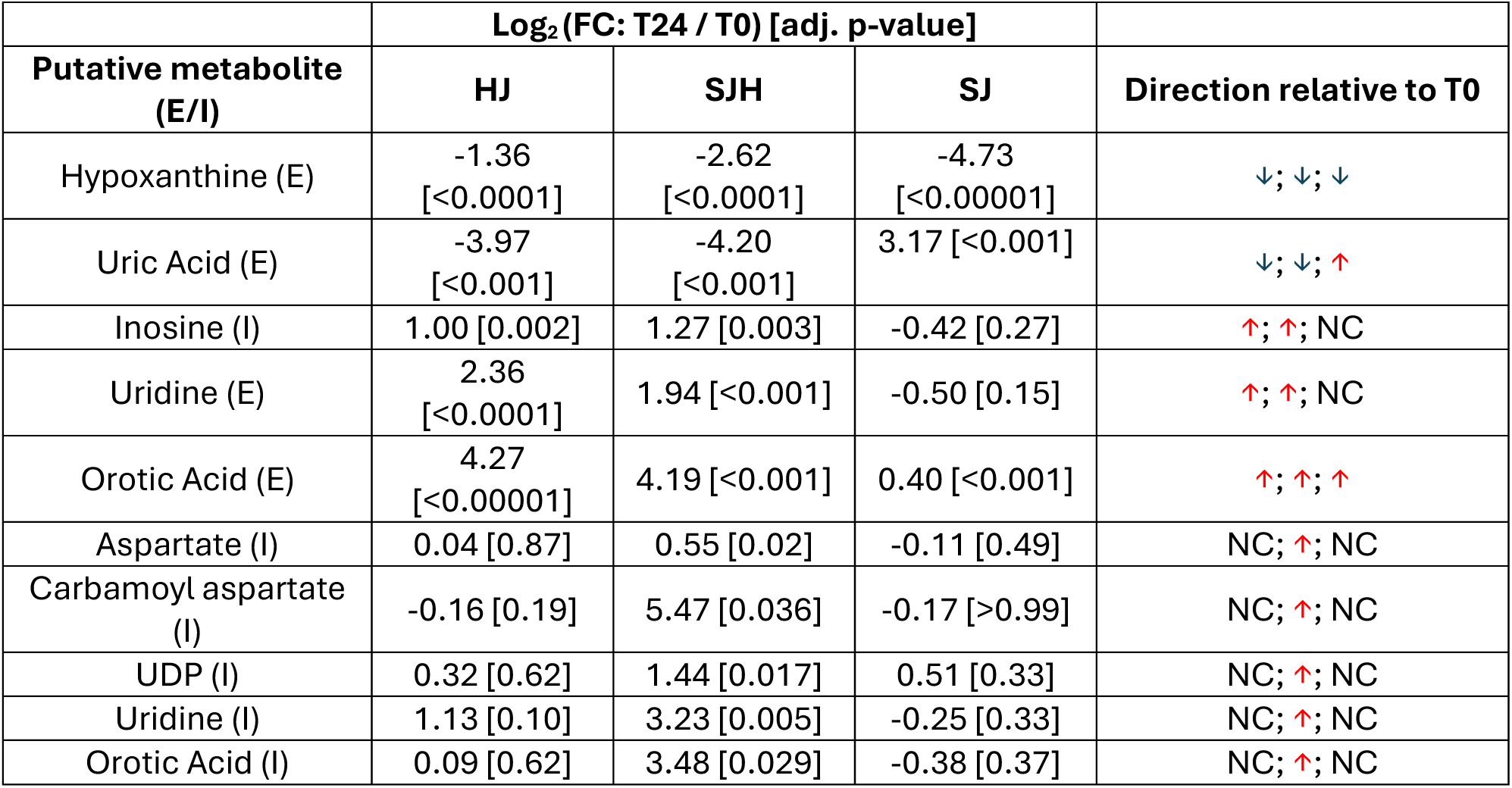
Fold change in extracellular and intracellular metabolite abundance after 24-hour incubation relative to starting media (t0). Adjusted p-value by t-test comparing T24 abundance to T0 abundance with BH correction for multiple testing. Abbreviations: FC: fold change; E: Extracellular; I: Intracellular; NC: no change.

#### Pyrimidine metabolism is increased in hepatocytes by co-culture with SW480 cells

Hepatocytes are a primary source of uridine, an intermediate of pyrimidine metabolism. We hypothesized that altered pyrimidine pools may be a result of newly available uridine from hepatocytes. Therefore, we measured pyrimidine intermediates by LC-MS/MS in media and cell extracts to observe changes in pool sizes over 24 hours compared to starting media. As expected, we observed an accumulation of uridine in the media of hepatocyte-containing co-cultures that was not present in SJ controls (**Figure 4D**). Interestingly, we observed an even greater accumulation of orotic acid, an intermediate of *de novo* pyrimidine biosynthesis (**Table 3**). We did not observe a significant difference between SJH and HJ controls, suggesting these media signals are primarily driven by hepatocytes. However, intracellular pool sizes of pyrimidine metabolites showed greater increases in SJH relative to HJ controls, including carbamoyl aspartate – the product of the rate-limiting step in *de novo* pyrimidine biosynthesis, suggesting the co-localization of hepatocytes and SW480s upregulates the pyrimidine biosynthetic pathway (**Figure 4E**).

### Discriminant ^13^C-ITUM analysis of SJH co-cultures

^13^C-stable isotope tracing of metabolic substrates has been used to detect metabolic adaptation (20, 28). However, incorporation of glucose-derived carbon in diverse metabolic pathways in mammalian cells convolutes biological interpretation of enrichment patterns in complex samples. We hypothesized that informatic integration of co-^13^C-enriched metabolites elucidates pathway activity. Therefore, we employed univariate and multivariate statistical approaches to characterize glucose utilization in co-culture. SJ and SJH co-cultures were treated with [U- ^13^C_6_]-glucose for 24 hours on day 9. On day 10, cells were snap frozen for ^13^C-ITUM and acquired LC-MS data was analyzed for ^13^C-enriched mass isotopomers (*i.e.,* isotopologues) [M+0, M+1, …, M+n] to identify nodes of glucose-derived metabolism that differentiate SJH co-cultures from SJ controls. We first used principal components analysis (PCA) to identify isotopologues that discriminate SJH co-cultures from HJ and SJ controls (**Figure 5A**). PC1 significantly separated SJH and SJ groups, while PC2 moderately separated SJH from HJ controls. Strong association with PC1 indicated co-^13^C-enrichment of these metabolites captures the impact of hepatocytes on SW480s. To further investigate adaptations to glucose metabolism after co-culturing hepatocytes on SW480s, we performed a univariate correlation analysis of top contributors to PC1 loadings, including only SJ and SJH ^13^C- enrichment. We visualized these relationships in a correlation matrix (**Figure 5B; Supplemental Figure 4**). A positive correlation between feature pairs indicates co-^13^C- enrichment of isotopologue pools in response to SW480 co-culture with hepatocytes relative to SW480 alone, while a negative correlation indicates a possible bifurcation of ^13^C- labeled carbon because of co-culture. As would be expected, fractional enrichment of M+0, the isotopologue indicating no ^13^C incorporation, of several metabolites, including TCA cycle intermediates, were positively associated with each other and clustered together (“Region 1” on **Figure 5B**). These M+0 species show a strong negative correlation with the ^13^C- enriched isotopologues (species in “Region 2”), *e.g.*, M+0 of Glutamate (“E_M0”) versus the incorporation of four ^13^C atoms (“E_M4”). This is the expected relationship and supports the validity of this analytical framework, which also reveals many unanticipated relationships. For example, the M+6 isotopologue of uridine diphosphate N-acetylglucosamine (UDPGlcNAc, corresponding to the direct incorporation of a labeled glucose molecule into glucosamine), clustered with unenriched (M+0) isotopologues of several metabolites in Region 1 (Red arrow, **Figure 5B**) and negatively correlated with enriched glutathione (*e.g.*, GSH_M3, found in Region 2; Blue arrow). UDPGlcNAc M+6 enrichment decreases in co-culture, while enrichment of GSH increases (relative to the SJ condition), suggesting glucose is redirected from the hexosamine biosynthetic pathway to glutathione synthesis in co-culture (**Supplemental Figure 5**). Region 3 shows a cluster of isotopologues from glycolytic and TCA cycle intermediates with strong co-enrichment. As expected, this includes enriched isotopologues of metabolites in the same pathway, such as glutamate (E_M4), a precursor to GSH (GSH_M4; Yellow arrows, **Figure 5B**). The positive co-enrichment of other TCA cycle intermediates with GSH may meet GSH demand in response to co-culture. Finally, Region 4 shows isotopologues with nominal relationships to each other, suggesting little change in response to co-culture. Together these data indicate ^13^C-enrichment of glutathione from glucose is significantly impacted by co-culturing of SW480s and hepatocytes. Glutathione is an important metabolite in redox homeostasis within the cell and changes to its biosynthesis may be a significant adaptation of metabolic pathways to the tumor-hepatocyte microenvironment.

**Figure 5.**
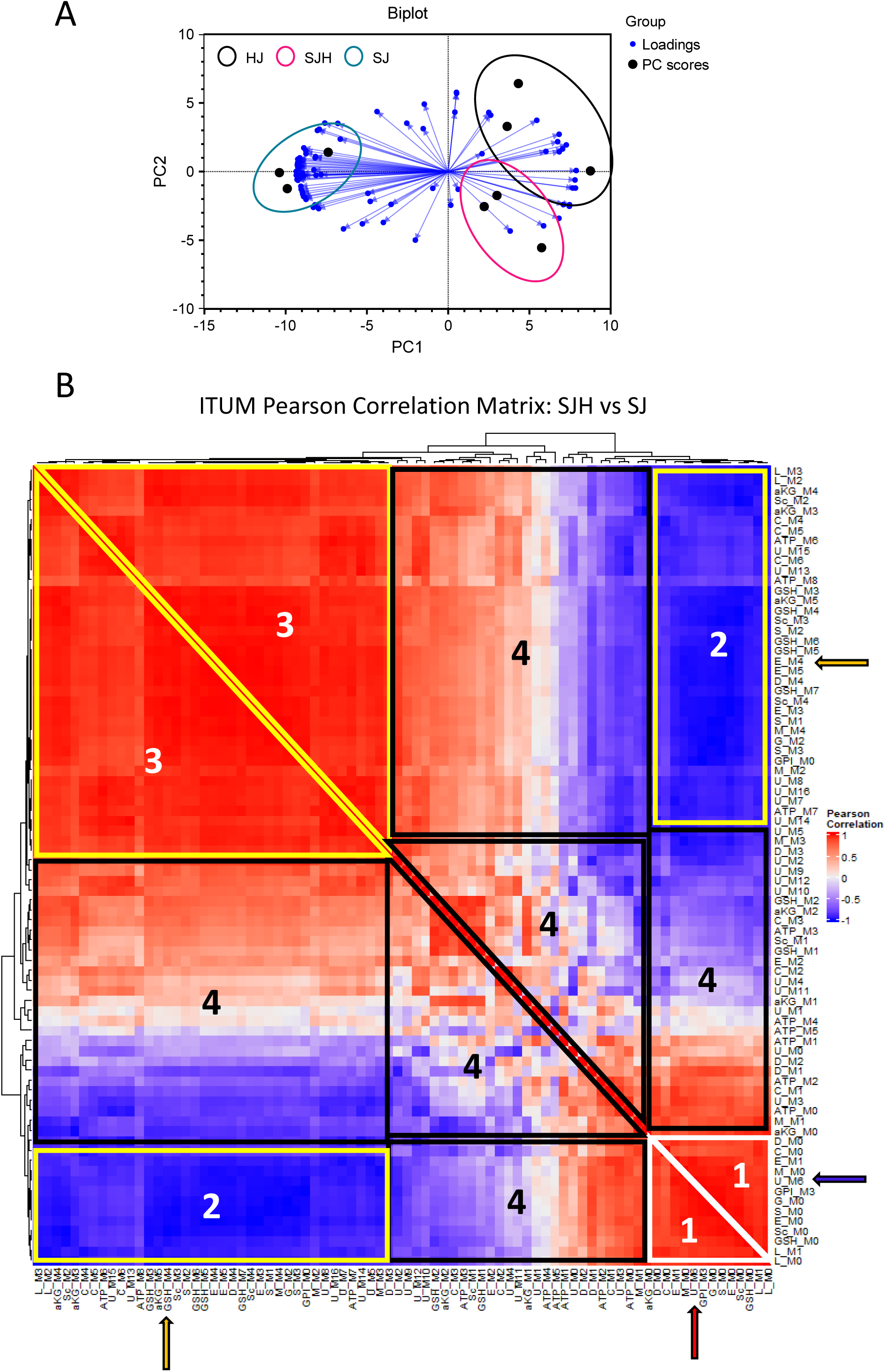
Discriminant ITUM analysis of SJH co-cultures. (A) Biplot of first two principal components (PC1, PC2) of PCA of HJ, SJH, and SJ co-cultures by ^13^C-glucose-enriched isotopologues. Black filled circles represent samples. Spheres show co-culture groups. Blue directed vectors show isotopologue loadings for PC1 and PC2. (B) Hierarchical clustering of ITUM SJH vs SJ Pearson correlation matrix. White triangle with “1” label indicates control cluster of isotopologues; Yellow shapes with “2” and “3” label corresponds to cluster of isotopologues from region of strongly co-enriched isotopologues in response to co-culture; Black shapes with “4” label corresponds to cluster of isotopologues with weak co-enrichment in response to co-culture. Red arrow, U_M6 positive correlation with unenriched M+0 isotopologues in Region 1; Blue arrow, U_M6 negative correlation with multiple isotopologues including GSH_M3 in Region 2; Yellow arrow, GSH_M4 positive correlation with metabolic precursor E_M4 in Region 3. Abbreviations: M#: isotopologue representing number of heavy carbons present in the molecule (*i.e.,* M1 indicates presence of 1 heavy 13-carbon), aKG: alpha-ketoglutarate, S: serine, M: Malate, L: lactate, C: citrate, ATP: adenosine triphosphate, U: uridine diphosphate N-Acetylglucosamine, G: glycine, D: Asparate, E: Glutamate, GSH: glutathione, Sc: Succinate, GPI: glycerophosphoinositol.

### Transcriptomic analysis of 3-dimensional hepatocyte-SW480 microtissue organoids

To evaluate SW480 adaptation to co-culturing with hepatocytes in a more physiological setting, we also performed transcriptomic profiling of SJ and SJH groups using a 3D microtissue organoid model. We co-cultured SW480 with primary rat hepatocytes and 3T3-J2 fibroblasts within collagen I-based microtissues (**Figure 6A**). Conditions included SJ cells and SJH using the same cell numbers and proportions as 2D co-culture. The SJH tricultures alongside co-culture controls were maintained for 7 days before harvesting for transcriptional and metabolic analyses. To identify transcriptional alterations that occur in cancer cells upon exposure to hepatocytes, we performed bead isolation using antibodies against CD326 (EpCAM) to isolate the SW480 tumor cells grown in the presence and absence of hepatocytes and supporting J2s and performed bulk RNA-seq analysis to identify genes and pathways that are modulated when exposed to hepatocytes. We found that exposure of tumor cells to hepatocytes led to increased expression of 708 genes and decreased expression of 762 genes in SW480 cells (adj. p <0.05) (**Figure 6B**). Further analysis of gene ontology (GO) and gene set enrichment analysis (GSEA) demonstrated significant alterations in hallmark pathways of Myc targets and pathways associated with several metabolic processes, including oxidative phosphorylation, metabolism of amino acids, GSH metabolism, and fatty acid metabolism (**Figure 6C-E; Supplemental Table 1,2**). We also compared RNAseq findings to oncogenic pathways (**Figure 6F**). These findings demonstrate that hepatocytes drive transcriptional changes in tumor cells, many of which are associated with changes in metabolic pathways.

**Figure 6.**
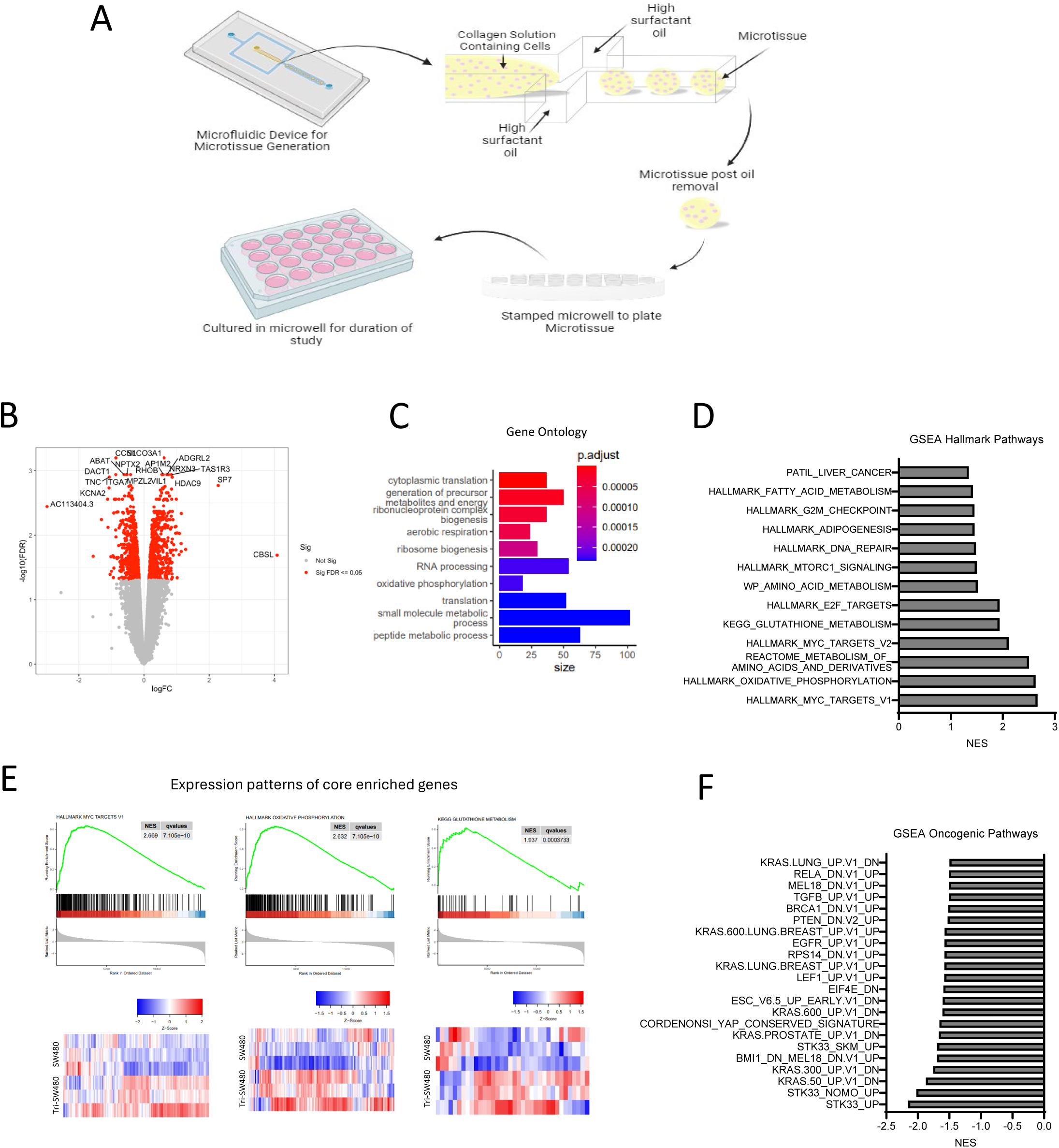
Transcriptional profiling of tumor cells in 3D microtissues identifies alterations in metabolic pathways upon exposure to hepatocytes. (A) 3D microtissue organoid scheme. (B) Volcano plot showing significantly upregulated and downregulated genes in samples from SJH cultures compared with SJ cultures. Positive logFC indicates up in SJH cultures. (C) Gene ontology analysis using DEG as input. (D) GSEA Hallmark analysis of SJ compared with SJH. A positive NES score indicates gene profiles that are enriched in tumor cells from the SJH condition compared with the SJ condition. All shown pathways adjusted FDR < 0.05. (E) Expression patterns of core enriched genes associated with Myc pathway and two metabolic pathways, oxidative phosphorylation and glutathione metabolism, that are positively enriched in the SJH condition and heat maps. (F) GSEA Oncogenic analysis of SJ compared with SJH. A negative NES score indicates gene profiles that are enriched in tumor cells from the SJ condition compared with the SJH condition. All shown pathways adjusted FDR < 0.05.

### Multiomics analysis of differentially expressed genes and static metabolite pools

Our 2D co-culture metabolomics pipeline and 3D transcriptional profiling both identified adaptations in amino acid, biosynthetic, and oxidative pathways. Therefore, we integrated these datasets in 2D to identify metabolite-gene relationships important to the SW480-hepatocyte microenvironment. Multiomic integration of a tumor cell transcription profile and metabolomic datasets can help establish the relationship between bulk metabolic adaptation and tumor cell phenotype. After bead isolation from hepatocytes and 3T3-J2s using antibodies against CD326 (EpCAM), RNA from SW480s was isolated and sequenced. Differentially expressed genes (adj. p < 0.01) were combined with significantly altered metabolite pools (**Figure 3B**) for univariate association analysis, recovering 627 mRNAs that were significantly correlated with 255 putative metabolites (p < 0.001, **Supplemental Table 3**). We further filtered this correlation matrix to only include very strong associations (R > |0.98|) to identify a subset of genes related to metabolic adaptation. A total of 151 unique genes correlated strongly with lactate, orotic acid, glutamyl-glycine (glu-gly), malate, and UMP (**Figure 7A**). In these analyses, positive correlations indicate co-expression in response to co-culture while negative correlations indicate opposing expression pattern in response to co-culture. Given our findings of glutathione metabolism, we chose to look closer at the 98 genes that associated with the metabolite glutamyl-glycine for gene-gene relationships (**Figure 7B**). GO enrichment analysis of biological processes of 98 genes associated with glu-gly revealed enrichment regulation of the cell cycle, response to hypoxia, and tissue morphogenesis (**Figure 7C**). Ontologies associated with all 151 genes further included adhesion, hypoxia, and angiogenesis (**Supplemental Figure 6A-B**).

**Figure 7.**
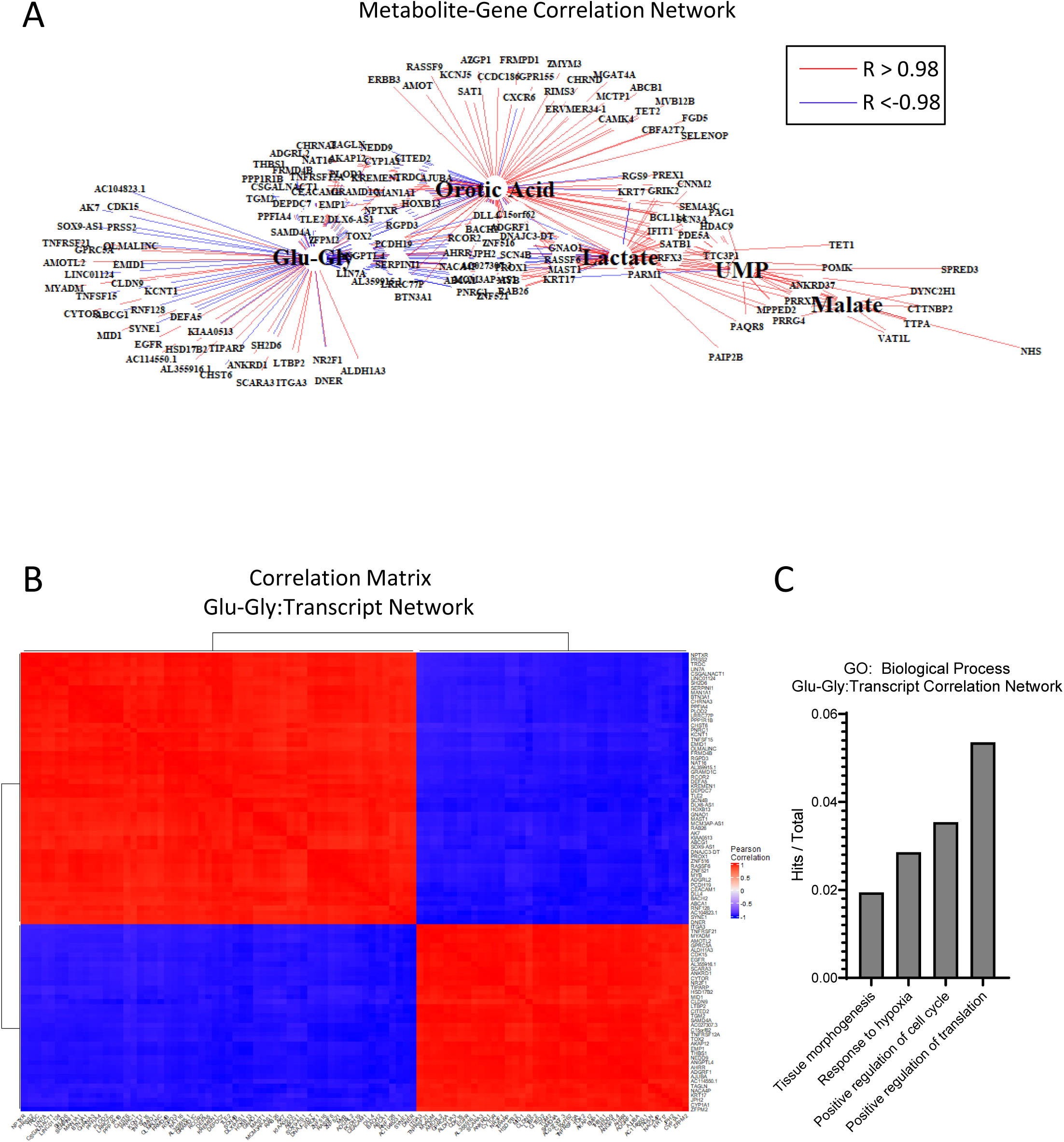
Multiomic pathway analysis of metabolic adaptation to hepatocytes. (A) Correlation network of differentially expressed genes (DEGs) and metabolites in SJH co-cultures compared to SJ control cultures. Gene names filtered from full bulk RNA-sequencing DEGs for significance of correlation to metabolites of interest (p < 0.001). Red lines indicate strong positive associations and blue represent strong negative associations (R >|0.98|). (B) Hierarchical clustering of Pearson correlation matrix of transcripts highly correlated with glutamyl-glycine (Glu-Gly). (C) Gene counts with functional group membership of 98 transcripts found to correlate strongly with glutamyl-glycine (Glu-Gly)

In this study we have demonstrated a multiomic platform that uses LC-MS-based untargeted metabolomics approaches and bulk RNA-sequencing to reveal complex interactions in a mixed cell population. Additionally, these approaches can be translated to 3D microtissue organoid models. Bioinformatic interrogation of datasets suggests an important adaptation in the oxidative environment and in reactive oxygen species (ROS) homeostasis in the tumor microenvironment upon introduction to hepatocytes.

## Discussion

This study employed a multiomic approach to uncover metabolic adaptation in the tumor-hepatocyte microenvironment using co-cultures of human colon adenocarcinoma cells (SW480) and primary rat hepatocytes, with the support of murine 3T3-J2 fibroblasts. By integrating untargeted metabolomics, ^13^C-stable isotope tracing untargeted metabolomics (ITUM), and transcriptomic analysis, we identified several critical metabolic alterations driven by the interaction between tumor cells and hepatocytes. First, co-culture of SW480s with primary hepatocytes significantly altered glucose metabolism. Hepatocytes reduced glucose consumption by 80% and lactate production by 94%, compared to SW480 monocultures. This suggests hepatocytes disrupt the classical Warburg effect in SW480 cells by influencing glucose metabolism (4, 27, 29). Previous work indicates that aerobic glycolysis is an important adaptation for overcoming hypoxia within the liver, thus suppression by hepatocytes may impact survival (11). Hepatocytes display highly dynamic metabolism that serves to regulate glucose homeostasis. In the presence of high glucose levels, hepatocytes can direct excess glucose molecules to storage as glycogen, *de novo* lipogenesis, or to produce uridine (30). Additionally, hepatocytes can use lactate for gluconeogenesis. Further study is necessary to deconvolute these intersectional and dynamic metabolic programs of varying glucose consumption (by both tumor cells and hepatocytes) and glucose production (hepatocytes) when cells are co-cultured in a manner that mimics the tumor microenvironment.

In response to reduced glucose consumption, SW480 cells adapted their fuel utilization. We observed a substantial reduction in ketone bodies and glutamine in the media of co-cultures, indicating increased consumption of these alternative substrates. In absence of glucose, tumor cells can reportedly shift to alternative fuels, such as glutamine, however whether this plasticity extends to ketone bodies is not yet well understood. Ketogenesis has been a topic of interest in cancer research (31). Studies have indicated a relationship between diminished expression of rate-limiting enzyme, 3-hydroxy-3-methylglutaryl synthase 2 (HMGCS2), and tumorigenesis (32–34). Thus, it is also possible diminished ketones in the media is due to reduced production through ketogenesis. Further, targeting ketogenesis has been proposed as a potential therapeutic strategy for inhibiting tumor progression, though responses *in vivo* vary between cancer types and stages (31, 35–38). Our studies suggest an impact of SW480-hepatocyte interaction on ketone body metabolism may represent adaptation for tumor survival. As ketogenic capacity has also been shown to vary across the spectrum of metabolic dysfunction-associated steatotic liver disease (MASLD), further study of the relationship between ketogenesis, cancer, and liver health is warranted.

Untargeted metabolomics revealed additional changes in metabolites associated with the TCA cycle and energy metabolism. Notably, acetyl-CoA was elevated, while pyruvate, ATP, and NAD^+^ were decreased, highlighting key metabolic nodes affected by hepatocyte presence. The co-culture system revealed a possible metabolic exchange between hepatocytes and SW480 cells, particularly in purine and pyrimidine metabolism. SW480s in co-culture displayed enhanced salvage and reduced uric acid production, coupled with increased pyrimidine biosynthesis. Uric acid can regulate oxidative stress, activity of enzymes related to glucose metabolism, and is associated with development of metabolic syndrome (39). Additionally, uric acid is linked with cancer risk (40, 41). Therefore, the modulation of uric acid levels and purine metabolism may be critical for targeting tumor growth. Further, due to comorbidities of cancer and obesity, the intersection of uric acid and metabolic syndrome in the tumor-hepatocyte niche may be an important area of further study. Recent work has established the pyrimidine intermediate, uridine, as an alternative fuel source for cancer cells in a glucose restricted environment (42, 43). As uridine was elevated in the media of hepatocyte-containing cultures, it is possible SW480s utilize this metabolite to drive activity in the pyrimidine pathway. Further study is needed to understand how access to uridine impacts cancer metabolism in the liver and how flux through these nucleoside pathways may impact the oxidative environment.

^13^C-stable isotope tracing of mixed cell populations results in convoluted ^13^C-enrichment datasets that can be difficult to interpret for individual cell populations. For this reason, studies often employ conditioned media exchange to study intercellular metabolic dependencies. These studies are limited by the lack of cellular contact that may be relevant to metabolic interactions and the difficulty adapting these methods toward the more complex and advantageous preclinical models of the microenvironment, such as 3D microtissue organoids. Using unsupervised dimension reduction and association analyses, discriminant ^13^C-ITUM allows for pathway analysis by identifying 1) enriched isotopologues that discriminate groups and 2) metabolite-metabolite relationships within the discriminating isotopologue set. Therefore, discriminant ^13^C-ITUM is a form of pathway analysis that highlights co-enriched metabolic pathways characterizing mixed cell populations. This systems-level approach enables specificity to substrate utilization with fewer constraints than metabolic flux analysis.

Transcriptomic profiling of SW480s revealed adaptation to metabolic pathways and enrichment of oncogenic pathways, including YAP, PTEN, and MYC. These signaling hubs have been implicated in the regulation of cancer cell metabolism (44–46). Here we observe their regulation in response to the presence of hepatocytes in the tumor microenvironment. Multiomic integration of metabolomics and RNA-seq data further showed that metabolic adaptations were associated with significant transcriptional changes in SW480 cells. These changes were linked to key biological processes, including adhesion, hypoxia response, and angiogenesis, indicating that tumor cells undergo functional adaptation in response to the hepatocyte microenvironment. These results are further supported by co-enrichment networks through a novel analysis of ITUM datasets, which we have called discriminant ITUM. This analysis indicated the importance of glutathione synthesis in differentiating SW480-hepatocyte co-cultures from SW480 controls. Finally, these findings were supported by preliminary studies of a 3D microtissue organoid model, which demonstrated that co-culturing SW480 cells with hepatocytes induces similar transcriptional and metabolic changes observed in 2D co-culture system.

While this study provides important insights into the metabolic interplay between tumor cells and hepatocytes, several limitations should be acknowledged. First, the use of 2D co-cultures, though effective in revealing key metabolic interactions, lacks the complexity of *in vivo* systems and may not fully capture the spatial and structural dynamics present in the liver microenvironment. Future work in advanced models such as organ-on-a-chip or 3D culture systems could provide more physiologically relevant insights. Additionally, our analysis primarily focused on the metabolic and transcriptional changes in tumor cells, leaving the metabolic impact on hepatocytes underexplored. Future work should involve a comprehensive assessment of hepatocyte responses to tumor interaction, including potential metabolic reprogramming. Mechanistic studies targeting the identified metabolic nodes, such as altered glucose and ketone metabolism or nucleoside exchange, could further elucidate their roles in tumor progression and present potential therapeutic strategies to disrupt these metabolic dependencies.

## Materials and Methods

### Reagents

LCMS grade water (H2O) (Fisher, W6-4), LCMS grade methanol (MeOH) (Fisher, A456-4), LCMS grade acetonitrile (ACN) (Fisher, A955-4), DMEM (high glucose) (Thermo, 11965092), DMEM (no glucose) (Thermo, A1443001), fetal bovine serum (Biotechne, S11150), Pen/Strep (Thermo, 15140122), L-glutamine (200 mM) (Thermo, 25030081), Phosphate Buffered Saline (PBS) (no CaCl2 or MgCl2) (Thermo, 14190144), Molecular Biology Grade Water (Corning, 46-000-CM), Pierce BCA Protein Assay Kit (Thermo, 23225), Genomic DNA kit (blood and cultured cells) (IBI scientific, IB47201). 0.25% Trypsin-EDTA (Corning 25-053-Cl), Hank’s Balanced Salt Solution (HBSS) (with CaCl2 or MgCl2) (Gibco 14025-076), Collagenase Type IV (Sigma Aldrich, C5138), Human CD326 (EpCAM) MicroBeads (Miltenyi Biotec, 130-061-101), CD16/CD32 Monoclonal Antibody (Invitrogen 14-0161-82), LS Magnetic Separation Columns (Miltenyi Biotec 130-042-401), QuadroMACS Magnetic Separator (Miltenyi Biotec 130-090-976), RNeasy Mini Kit (Qiagen, 74104).

### 2D and 3D culture platform

For both 2D and 3D cultures, 3T3-J2 fibroblasts (Kerafast, Catalog Number: EF3003) and SW480 cells (ATCC, lot Number: 700031955) are maintained in tissue culture flasks until ready for use. Primary rat hepatocytes (PRH, Cryopreserved Male Wistar Rat Plateable Hepatocytes AMY 7 mil, Catalog Number: r3000.H15 Lot No. 1210326) were thawed immediately before use. 3T3-J2 fibroblasts are growth arrested using 1 µg/mL mitomycin-C for 4 hours in culture before detachment using trypsin-EDTA.

In 2D, 150k PRH and 150k 3T3-J2 fibroblasts are plated per well for all HJ, SJH, SJ wells. On day 7, 50k SW480 cells are seeded to SJH and SJ wells. On day 9, cell culture media is changed to exchange equimolar (22 mM) unlabeled glucose for non-radioactive, stable isotopically labeled 22 mM [U-^13^C_6_]glucose according to previously established protocols (47). Cells and final conditioned media are collected on day 10.

3D microtissue organoids were fabricated as previously described (25). Briefly, plates were with 2% agarose and allowed to stiffen for 24 hours. We then used a microwell stamp made from a PDMS mold to create 200 μM microwells that held the microtissues separately within the same well. After cleaning the microwells with a series of washes, we added media to the wells. We maintained the microtissues in Human Hepatocyte Maintenance Media (HHM) (43.25mL 1x DMEM, 5 mL Bovine Serum, 750 μL HEPES, 1M, pH 7.6, 500 μL insulin/human transferring/selenous acid and linoleic acid premix (Corning premix solution), 500 μL Penicillin-streptomycin, 100X solution of 50 mg/mL stock, 0.5 μL Dexamethasone, 10 mM in DMSO, 0.5 μL Glucagon, 0.7 mg/mL in 0.05 M acetic acid. During the study, we provided fresh media changes every 48 hours by removing 300 μL from each well and replacing it with 300 μL of fresh media. We then removed the microtissues at their respective time point.

### Quantification of Glucose and Lactate via 1 H NMR

∼50 µL of cell culture media was dried to completion in a SpeedVac in the presence of D_2_O to aid in water suppression. Samples were reconstituted in D_2_O (99.9%) spiked with 0.3 mM d_4_-trimethyl-silyl propionate (TSP). ^1^H-NMR signals were acquired using a Bruker Avance III 600 NMR instrument equipped with a CryoProbe, then the integrated intensities of the α-anomeric proton on glucose carbon-1, the methyl signal for lactate, and the tri-methyl signal from TSP, were used to calculate molar concentrations of the respective substrates. For all ^1^H-NMR collections, spectra were collected by conventional pulse-and-collect measurements under quantitative conditions (10-ppm spectral range using ∼15 μs [90°] excitation pulse and 22-second delay between each of 20 transients).

### Untargeted Metabolomics and Isotope Tracing Untargeted Metabolomics pipeline

Cells are harvested, metabolites extracted, and raw data acquired using liquid chromatography (LC) on a Thermo Vanquish UHPLC system, and full-scan high-resolution mass spectrometry (HR-MS) on a Thermo QExactive plus hybrid quadrupole-orbitrap mass spectrometer fitted with heated electrospray ionization source and operated in negative and positive polarity mode.

#### Cell collection for metabolomics

Media was collected and snap frozen, then adherent cells were washed twice with 1 mL warm (37°C) PBS (-MgCl2, -CaCl2), once with warm (37°C) cell-culture grade H2O, then the entire plate was submerged into liquid nitrogen to snap freeze cells, which rapidly quenches metabolism. To preserve the metabolome, cells were scraped in 500 μL of cold (-20°C) LCMS grade MeOH per well of cells. Two wells were combined to form each replicate, n = 3 per group. Finally, MeOH was evaporated using a SpeedVac. Dried cell pellets were stored at -80°C until analysis.

#### Metabolite extraction

Metabolites from cell pellets and conditioned media were extracted and analyzed by LC-MS untargeted metabolomics according to previously published protocols (47, 48). Briefly, cell pellets are reconstituted in 1 mL 2:2:1 (v/v/v) ACN:MeOH:Water, then vortexed (30s), flash frozen in liquid nitrogen (1 min), and sonicated (25°C, 10 min) in three cycles. After 1 hour at -20°C, samples are centrifuged at 15k x g at 4°C for 10 minutes. Supernatant was transferred to a fresh tube and dried by SpeedVac overnight, while the remaining cell pellet was stored at -80°C after removing any remaining solvent for DNA quantification. Cell extracts are reconstituted in 40 µL 1:1 (v/v) ACN:Water for analysis. 20 µL of conditioned media was extracted in 80 µL 1:1 (v/v) ACN:MeOH. Media underwent a single cycle of vortex and sonication, then followed the same procedure as cell pellets. Media samples were reconstituted in 200 µL 1:1 (v/v) ACN:Water for analysis.

#### Data acquisition

For this study, polar metabolites were acquired using hydrophilic interaction chromatography (HILIC) and energy nucleotides were acquired by reverse phase (RP), using two unique UHPLC methods: **[1]** SeQuant ZIC-pHILIC column (2.1 x 150 mm, 5 μm) (Millipore Sigma, 1.50460). Mobile phase A (MPA) was 95% H2O, 5% ACN, 10 mM ammonium acetate, and 10 mM ammonium hydroxide. Mobile phase B (MPB) was 100% ACN. The total run time was 50 minutes, flow rate was 2 mL/min, column chamber was set to 45°C, and 2 µL sample was injected. Mobile phase gradient was as follows: 0-0.5 min, 90% MPB; 0.5-30 min, 90→30% MPB; 30-31 min, 30% MPB; 31-32 min, 30→0% MPB; 32-33 min, 0→90% MPB; 33-50 min, 90% MPB. **[2]** Energy nucleotides (ATP, ADP and AMP), and redox nucleotides (NAD^+^ and NADH) were measured as previously described, with modifications. Briefly, metabolites were extracted from cells for ITUM in MeOH:ACN:H2O (2:2:1), then extracts were separated and detected using ion-pairing RP UHPLC-MS/MS on a C18 column (Waters Xbridge, 150 x 2.1mm, 3μm). Nucleotides were detected as adducts of dibutylamine acetate on a Thermo QExactive Plus mass spectrometer, operated in positive ionization mode, using parallel reaction monitoring transitions as previously described.

For all metabolomics pipelines, both blanks and pooled quality control (QC) samples are injected periodically throughout the run. Blanks were ACN:H2O (1:1) and the QC sample was a pooled sample including all naturally-occurring and ^13^C-labeled samples. To aid in chemical feature identification, two additional samples were also injected. First, was a standard mix consisting of authentic standards, for all expected analytes. Second, was a pooled sample consisting only of the naturally-occurring samples, analyzed via data-dependent analysis (DDA) tandem mass spectrometry (MS/MS) using IE omics script and R-Studio (49).

#### Data preparation

Data processing and initial analysis was performed using Thermo Compound Discoverer 3.3. After raw mass spectra were uploaded, background ions were removed, retention time (RT) for detected signals were aligned across samples, chemical formulas were predicted, then grouped chemical features were profiled to determine compound identity, based on (1) the m/z predicted from the chemical formula, (2) the RT compared to an authentic external standard, and (3) the MS/MS fragmentation pattern, compared to in-house standards or online databases.

For ITUM experiments, putatively identified metabolites and lipids were then carried forward and [^13^C] stable-isotope enrichment with correction for natural abundance. To carry out [^13^C] stable isotope tracing, all [^13^C] mass isotopomers (*i.e.,* isotopologues) within the isotopic envelope of each identified metabolic pool were identified based on the diagnostic shift in m/z (Δm/z = 1.0033 Da, natural abundance, 1.11% of all carbon) induced by the presence of ^13^C-labeled compounds. Raw ion counts for each isotopologue were extracted, summed, and expressed as a percentage of the total pool. After natural abundance correction, the fractional intensities were then graphed as a function of [^13^C] content, generating mass isotopologue distributions (MIDs) for each detected metabolite or lipid.

For static pool analysis, total ion counts were exported from Compound Discoverer and normalized to biomass (either total DNA or total protein). DNA was quantified from cell pellets after metabolite extraction using IBI Scientific genomic DNA kit for cultured cells as previously described for metabolite normalization following kit instructions (50). Total protein was quantified using the Pierce BCA Protein Assay Kit from separately cultured and harvested samples in parallel with metabolomics experiments.

### Quantification of Total Ketone Bodies

Acetoacetate (AcAc) and β-hydroxybutyrate (BOHB) were formally quantified using UHPLC-MS/MS as described previously (51, 52). Briefly, [U-^13^C_4_]AcAc and [3,4,4,4-D4]βOHB internal standards were spiked into ice cold MeOH:ACN (1:1), then ketones were extracted, separated via reverse-phase UHPLC, and detected via parallel reaction monitoring (PRM) on a QExactive Plus hybrid quadrupole-orbitrap mass spectrometer.

### Bulk RNA sequencing and analysis

For the 2D platform, cells were collected from the plates using 0.25% Trypsin-EDTA. For the 3D platform, microtissues were collected from the wells and dissociated by incubating at 37°C with 0.1mg/mL Collagenase Type IV in HBSS (+CaCl2, +MgCl2) for 30 minutes, then manually disrupted to form a single cell suspension. For both platforms, 8 wells were combined to form replicates, n=3 per group. The single cell suspensions were incubated with MicroBeads against human CD326 (EpCAM) and antibody against CD16/CD32 to block nonspecific binding, then isolated across Miltenyi LS magnetic separation columns. RNA was isolated from the sorted SW480 cells using a Qiagen RNeasy Mini Kit.

#### Bulk RNA-seq Analysis

RNA-seq analysis was performed at the Minnesota Supercomputing Institute at the University of Minnesota. Briefly, Fastq files were first processed with the CHURP pipeline (version 0.2.3) (53) to perform adaptor trimming using trimmomatic (version 0.33) (54); reads were then mapped to Homo sapiens GRCh38 genome using HiSat2 (version 2.1.0) (55). Subreads count was generated using the Subreads featureCounts tool (version 1.6.2) (56) using the Homo_sapiens.GRCh38.100.gtf annotation. Count data were filtered by removing genes that were less than 300 nt in length and including only genes that had a cpm (counts per million) value greater than 1 cpm in at least two sample replicates. The quasi-likelihood test was used to evaluate differential expression (DE) with edgeR (version 3.38.1) (57, 58). The Benjamini-Hochberg method was used to adjust p-values for multiple hypothesis testing and an adjusted p-value ≤ 0.05, with a log2 fold change > 0 was used as a DE significance threshold. For gene ontology (GO pathway and GSEA analysis, the R package clusterProfiler (version 4.4.4) (59, 60) was used. Normalized Enrichment Score (NES) indicates the distribution of genes across a ranked list and normalizes the correlation between gene sets and datasets to gene set size, allowing for comparison. For GSEA analysis specifically, Hallmark and 18 selected C2 pathways were combined and used for testing (see full GSEA pathway results and selected C2 pathways in **Supplemental Table 1, 2**). Analysis of GSEA C6 oncogenic pathways, read counts were converted to normalized counts using the DESeq2 (version 1.42.0) R package and were further filtered to get rid of genes with zero expression across all samples. GSEA was then performed using the standalone software (version 4.3.2) developed by the Broad Institute, and statistical significance was assessed via gene set permutation testing (1000 permutations) (61, 62). GEO accession number GSE282081.

### Statistics and Multi-omic Analysis

Descriptive data are expressed as mean and standard error (SEM) for continuous measures. Comparison of metabolite abundance were made between HJ and SJH or SJ and SJH by unpaired t-test and corrected for multiple comparisons by Benjamini-Hochberg using GraphPrism v10.2.3. Multi-omic integration of bulk transcriptomic and metabolomics datasets were performed in R v4.2.3 using rcorr() from the Hmisc package and the igraph package for visualization of correlations. Given the small dataset, only highly significant correlations (p-value of Pearson correlation coefficient < 0.001) were included in downstream pathway and gene ontology analysis. Genes that correlated strongly with metabolites of interest were included as a set of gene IDs and fold changes in an ExpressAnalyst query for functional analysis (63).

Discriminant ITUM was performed using a curated dataset of positively identified metabolites by commercial standard and / or MS/MS match in mzCloud. Selected metabolites showed significant total enrichment in the SJH group after correction for natural abundance of ^13^C (1.11% of all carbon in nature), performed within the Compound Discoverer “stable isotope labeling” node based on predicted chemical formula. All isotopologues with enrichment were included for a given metabolite as individual variables, *i.e.* S_M0, S_M1, S_M2, S_M3 included as four unique variables representing serine enrichment. Data presented in 0-100 range and represents percentage of the total metabolite pool detected by LC-MS in full scan (MS1). Total pools are calculated by sum of ion counts for each possible isotopologue (*i.e.* total ion counts of serine = S_M0 + S_M1 + S_M2 + S_M3; % Enrichment of S_M3 = ion counts of S_M3 / total ion counts of serine * 100). The resulting multivariate dataset was used for Principal Components Analysis (PCA) to identify co-enriched isotopologues that discriminate SJH co-cultures from SJ or HJ controls. PCA was performed in GraphPrism v10.2.3. Data was standardized prior to PCA. A correlation network was graphed from the top contributing isotopologues distinguishing SJH from SJ controls using the igraph package in the R environment for qualitative pathway analysis of glucose utilization in mixed cell populations. Joint pathway analysis of 3D microtissue transcriptomics and metabolomics datasets performed in MetaboAnalyst 4.0 (64).

## Acknowledgements

We are grateful to Alisha Seay for technical support. This study was funded by the University of Minnesota, Medical School Dean’s Academic Investment Research Program (AIRP). Additional funding: R01DK091538 (PAC); T32DK007203 (ABN); R01CA215052 (KLS); HL166142 (EDQ).

## Figure Legends

**Supplemental Figure 1.**
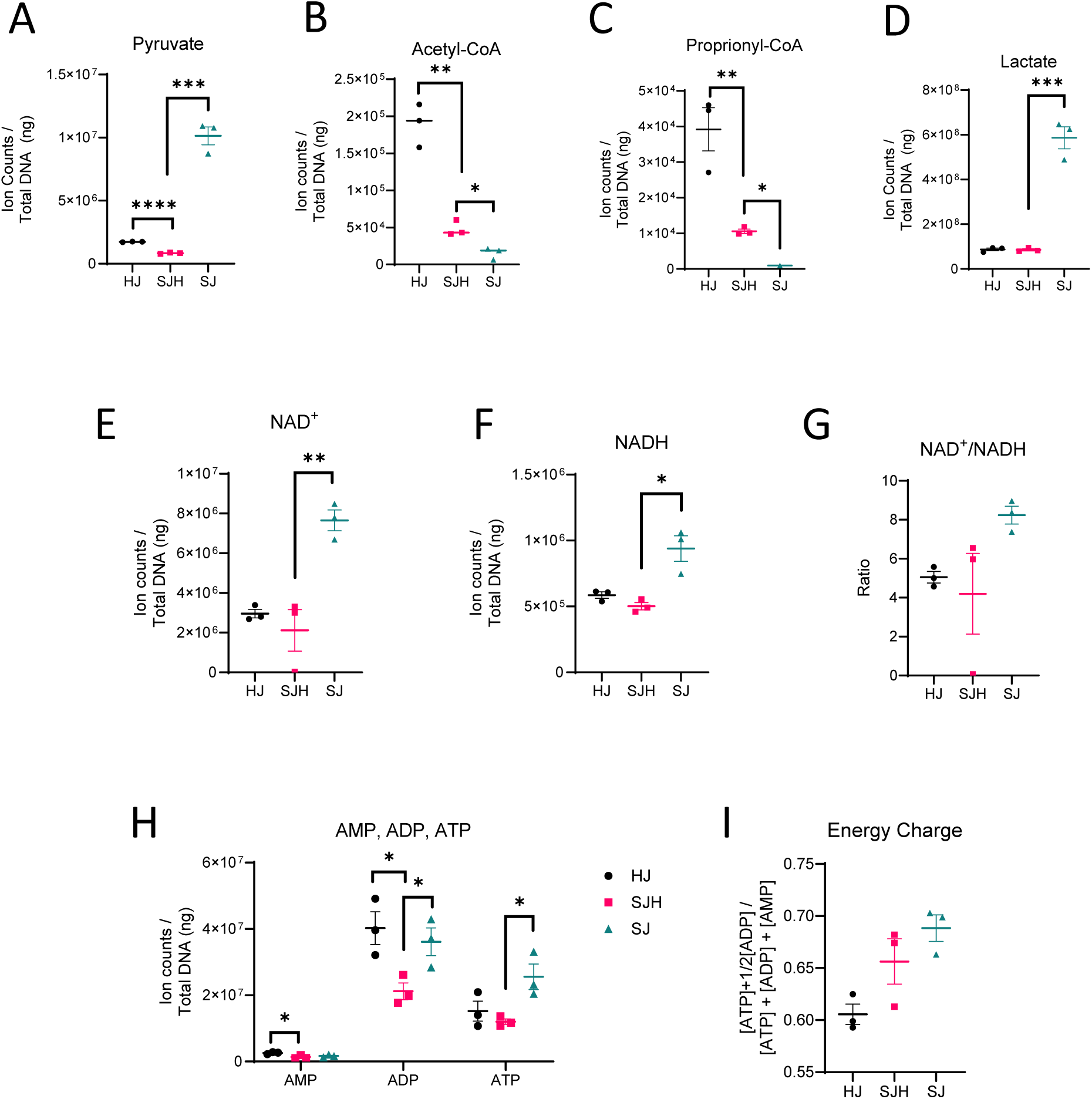
Metabolite abundance in 2D co-cultures. Ion counts after normalization to total ng DNA of: (A) pyruvate, (B) Acetyl-CoA, (C) Propionyl-CoA, (D) Lactate, (E) NAD^+^, (F) NADH, and (H) AMP, ADP, ATP nucleotides. (G) Ratio of NAD^+^ to NADH ion counts. (I) Calculated energy charge of each co-culture group based on ion counts of AMP, ADP, and ATP. Significance tested using unpaired t-test, comparison HJ vs SJH or SJ vs SJH, and corrected for multiple comparisons using Benjamini-Hochberg method. *: p adj. < 0.05, **: p adj. < 0.01, ***: p adj. < 0.001, ****: p adj. < 0.0001.

**Supplemental Figure 2.**
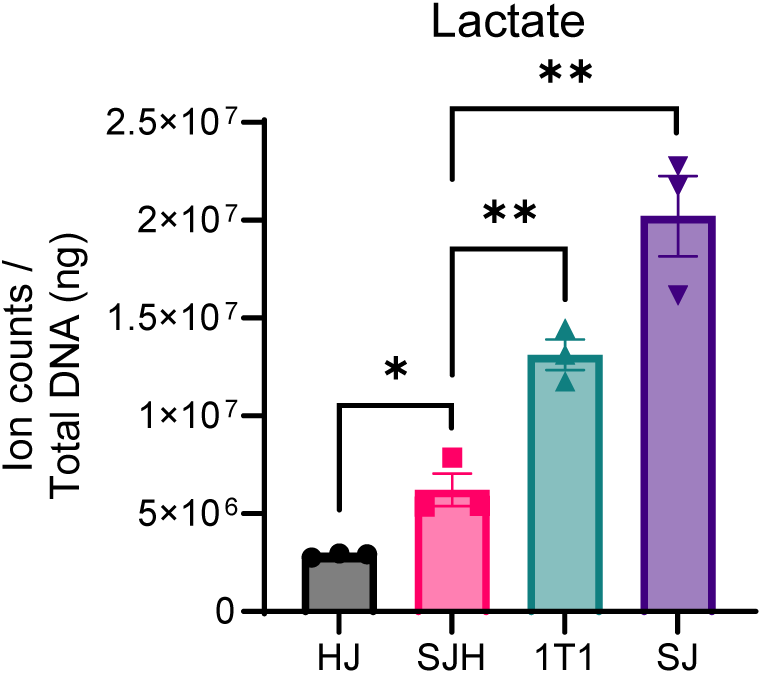
Lactate abundance in co-culture and 1T1 dilution. Bar graph of total ion counts after normalization to DNA compared to analytical dilution (1T1). Significance tested using unpaired t-test, comparison HJ vs SJH or SJ vs SJH, and 1T1 vs SJH; corrected for multiple comparisons using Benjamini-Hochberg method. *: p adj. < 0.05, **: p adj. < 0.01

**Supplemental Figure 3.**
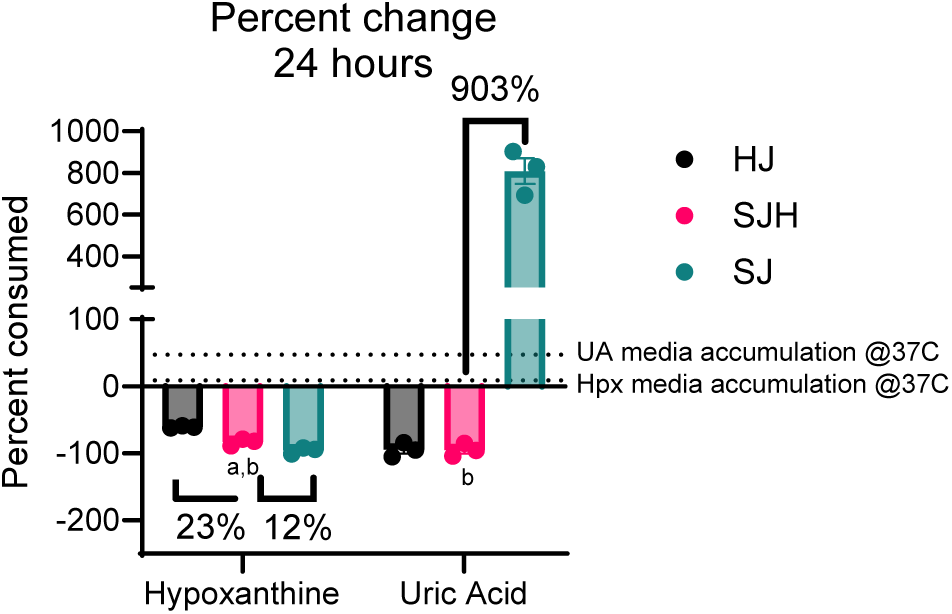
Percent change in media purines. Percent difference in media after 24h incubation with co-cultured cells where time point 0 represents starting media abundance prior to cell exposure. Dotted lines represent accumulation of hypoxanthine (Hpx) and uric acid (UA) after 24h at 37°C in media in absence of cells. Statistical comparison by unpaired t-test; letters indicate significance in comparison to HJ controls (“a”) or SJ controls (“b”).

**Supplemental Figure 4.**
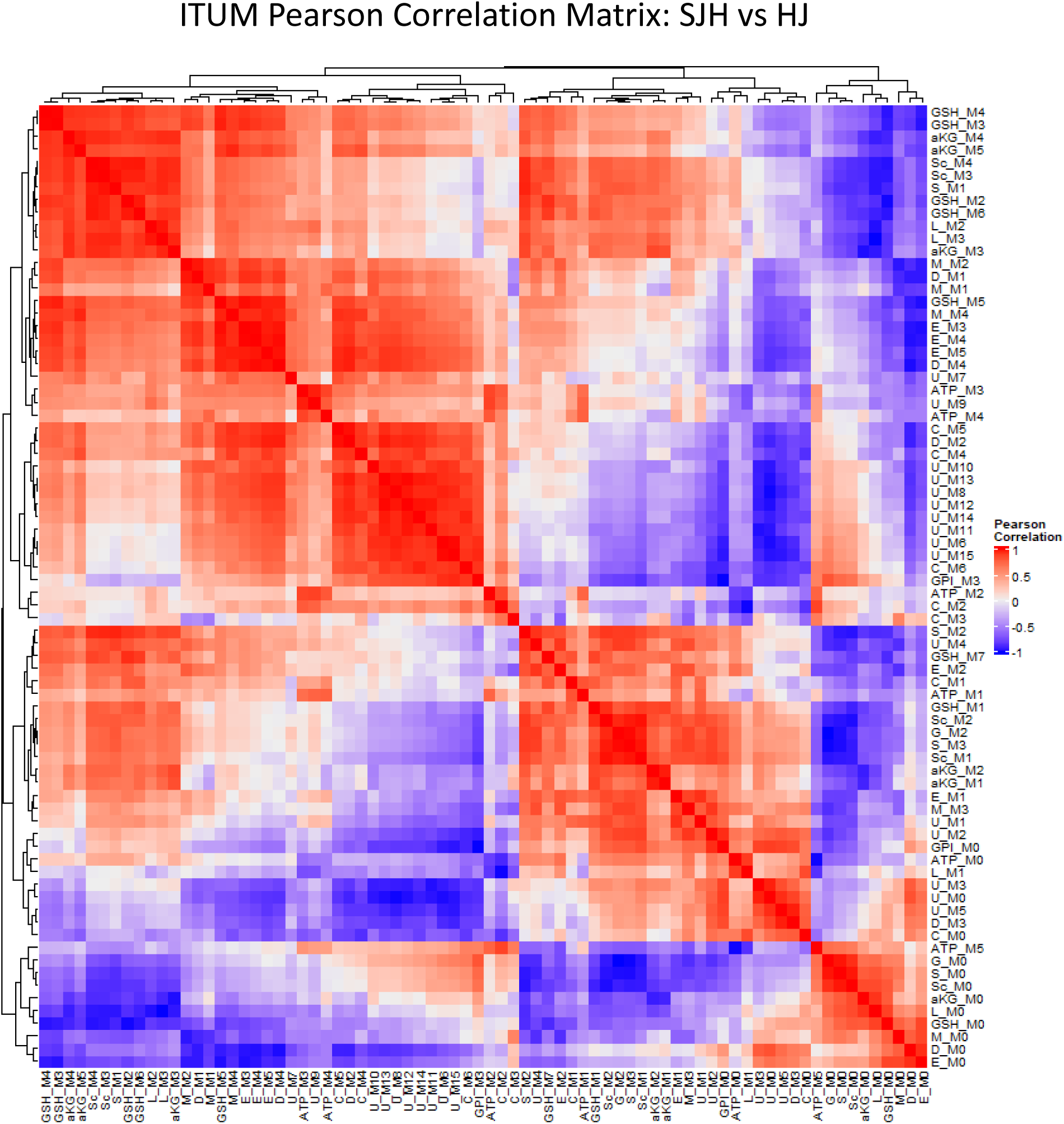
Correlation matrix of SJH vs HJ ITUM in 2D co-cultures. Correlation matrix assessing co-enriched isotopologues in response to presence of hepatocytes (SJ, SJH cultures). Red gradient represents positive associations while blue represents negative associations. Correlations measuring by Pearson correlation method.

**Supplemental Figure 5.**
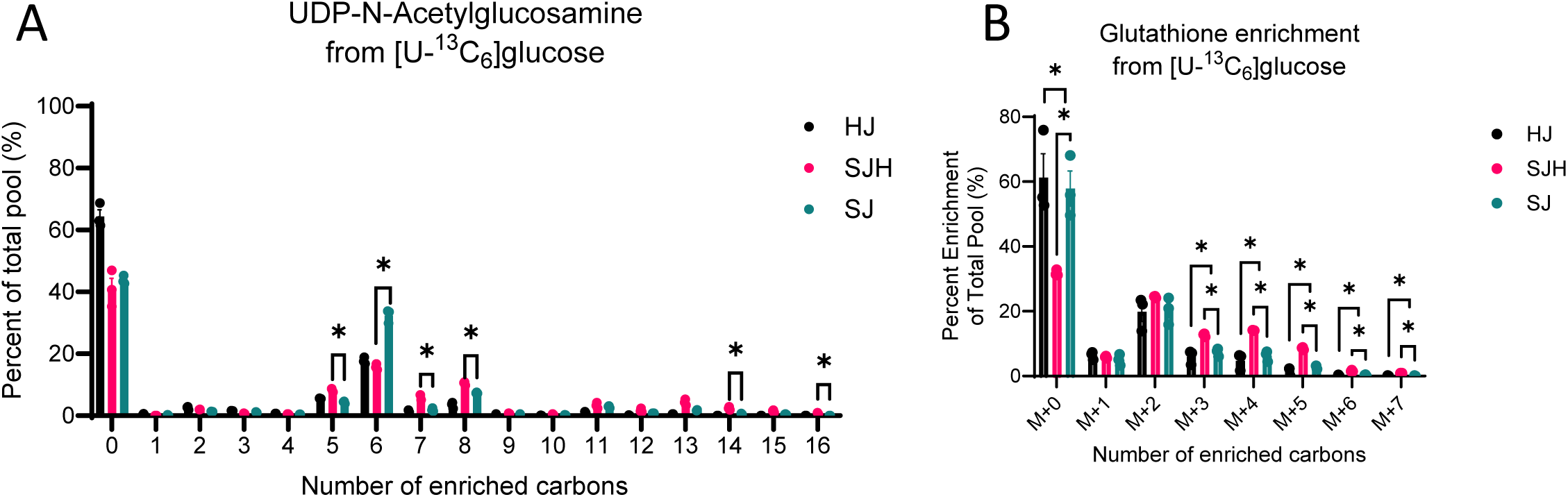
Distribution of ^13^C enrichment. Percent enrichment of total pools of (A) Uridine diphosphate N-acetyglucosamine and (B) glutathione after 24h incubation with [U-^13^C_6_]glucose at 37°C. *: p adj. < 0.05

**Supplemental Figure 6.**
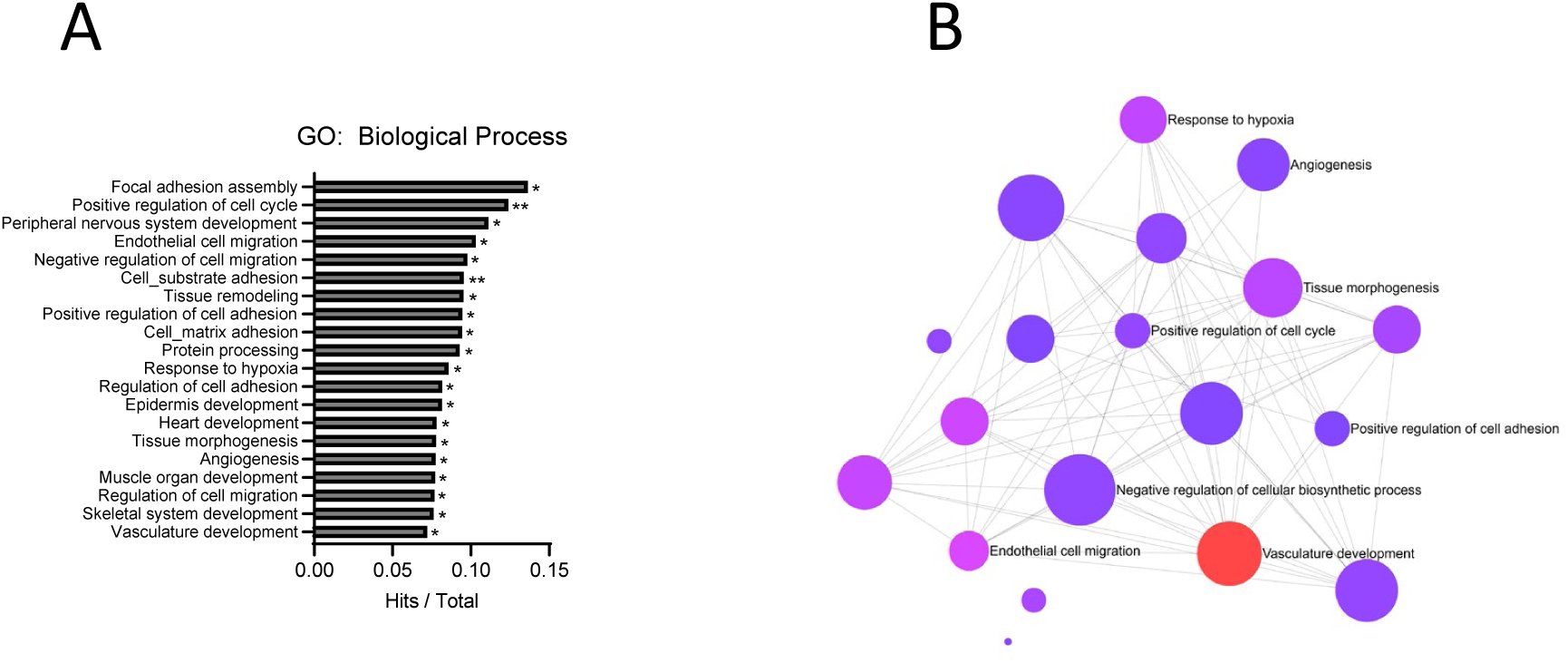
Multiomic pathway analysis. (A) Gene counts with functional group membership of 151 genes found to correlate strongly with glutamyl-glycine (Glu-Gly), orotic acid, lactate, uridine monophosphate, and malate. (B) Functional network of shared genes in represented transcriptional profile with strong metabolite-gene associations; analysis performed using ExpressAnalyst. (C) Scatter plot of joint pathway analysis from MetaboAnalyst v4.0 using DEGs and full static metabolomics dataset based on associated fold changes. X and y axes show enrichment score in genes and metabolite peaks, respectively.

## Notes

https://www.ncbi.nlm.nih.gov/geo/query/acc.cgi?acc=GSE282081

### Competing Interest Statement

P.A.C. has served as an external consultant for Pfizer, Inc., Abbott Laboratories, Janssen Research & Development and Juvenescence.

### Summary of Updates

This version includes minor improvements in descriptions and interpretations. No new data compared to the first submission are enclosed.

## References

1. Siegel RL, Giaquinto AN, and Jemal A. Cancer statistics, 2024. CA: A Cancer Journal for Clinicians. 2024;74(1):12–49.

2. Horn SR, Stoltzfus KC, Lehrer EJ, Dawson LA, Tchelebi L, Gusani NJ, et al. Epidemiology of liver metastases. Cancer Epidemiology. 2020;67:101760.

3. Hanahan D, and Weinberg Robert A. Hallmarks of Cancer: The Next Generation. Cell. 2011;144(5):646–74.

4. Pavlova Natalya N, and Thompson Craig B. The Emerging Hallmarks of Cancer Metabolism. Cell Metabolism. 2016;23(1):27–47.

5. Lyssiotis CA, and Kimmelman AC. Metabolic Interactions in the Tumor Microenvironment. Trends in Cell Biology. 2017;27(11):863–75.

6. Sullivan MR, Danai LV, Lewis CA, Chan SH, Gui DY, Kunchok T, et al. Quantification of microenvironmental metabolites in murine cancers reveals determinants of tumor nutrient availability. eLife. 2019;8:e44235.

7. Lin J, Rao D, Zhang M, and Gao Q. Metabolic reprogramming in the tumor microenvironment of liver cancer. Journal of Hematology & Oncology. 2024;17(1):6.

8. Fowle-Grider R, Rowles JL, Shen I, Wang Y, Schwaiger-Haber M, Dunham AJ, et al. Dietary fructose enhances tumour growth indirectly via interorgan lipid transfer. Nature. 2024.

9. Huang Q, Tan Y, Yin P, Ye G, Gao P, Lu X, et al. Metabolic Characterization of Hepatocellular Carcinoma Using Nontargeted Tissue Metabolomics. Cancer Research. 2013;73(16):4992–5002.

10. Nwosu ZC, Megger DA, Hammad S, Sitek B, Roessler S, Ebert MP, et al. Identification of the Consistently Altered Metabolic Targets in Human Hepatocellular Carcinoma. Cellular and Molecular Gastroenterology and Hepatology. 2017;4(2):303–23.e1.

11. Dupuy F, Tabariès S, Andrzejewski S, Dong Z, Blagih J, Annis Matthew G, et al. PDK1-Dependent Metabolic Reprogramming Dictates Metastatic Potential in Breast Cancer. Cell Metabolism. 2015;22(4):577–89.

12. Bu P, Chen K-Y, Xiang K, Johnson C, Crown SB, Rakhilin N, et al. Aldolase B-Mediated Fructose Metabolism Drives Metabolic Reprogramming of Colon Cancer Liver Metastasis. Cell Metabolism. 2018;27(6):1249–62.e4.

13. Loo Jia M, Scherl A, Nguyen A, Man Fung Y, Weinberg E, Zeng Z, et al. Extracellular Metabolic Energetics Can Promote Cancer Progression. Cell. 2015;160(3):393–406.

14. Clish CB. Metabolomics: an emerging but powerful tool for precision medicine. Cold Spring Harb Mol Case Stud. 2015;1(1):a000588.

15. Patti GJ, Yanes O, and Siuzdak G. Innovation: Metabolomics: the apogee of the omics trilogy. Nat Rev Mol Cell Biol. 2012;13(4):263–9.

16. Nelson AB, Chow LS, Stagg DB, Gillingham JR, Evans MD, Pan M, et al. Acute aerobic exercise reveals FAHFAs distinguish the metabolomes of overweight and normal weight runners. JCI Insight. 2022.

17. Jang C, Chen L, and Rabinowitz JD. Metabolomics and Isotope Tracing. Cell. 2018;173(4):822–37.

18. Puchalska P, Martin SE, Huang X, Lengfeld JE, Daniel B, Graham MJ, et al. Hepatocyte-Macrophage Acetoacetate Shuttle Protects against Tissue Fibrosis. Cell Metab. 2019;29(2):383–98 e7.

19. Stagg DB, Gillingham JR, Nelson AB, Lengfeld JE, d’Avignon DA, Puchalska P, et al. Diminished ketone interconversion, hepatic TCA cycle flux, and glucose production in D-β-hydroxybutyrate dehydrogenase hepatocyte-deficient mice. Molecular Metabolism. 2021;53:101269.

20. Puchalska P, Huang X, Martin SE, Han X, Patti GJ, and Crawford PA. Isotope Tracing Untargeted Metabolomics Reveals Macrophage Polarization-State-Specific Metabolic Coordination across Intracellular Compartments. iScience. 2018;9:298–313.

21. Buescher JM, Antoniewicz MR, Boros LG, Burgess SC, Brunengraber H, Clish CB, et al. A roadmap for interpreting 13C metabolite labeling patterns from cells. Current Opinion in Biotechnology. 2015;34:189–201.

22. Muir A, Danai LV, and Vander Heiden MG. Microenvironmental regulation of cancer cell metabolism: implications for experimental design and translational studies. Disease Models & Mechanisms. 2018;11(8):dmm035758.

23. Xiang C, Du Y, Meng G, Soon Yi L, Sun S, Song N, et al. Long-term functional maintenance of primary human hepatocytes in vitro. Science. 2019;364(6438):399–402.

24. Sun P, Zhang G, Su X, Jin C, Yu B, Yu X, et al. Cell Reports. 2019;29(10):3212–22.e4.

25. Kukla DA, Crampton AL, Wood DK, and Khetani SR. Microscale Collagen and Fibroblast Interactions Enhance Primary Human Hepatocyte Functions in Three-Dimensional Models. Gene expression. 2020;20(1):1–18.

26. DeBerardinis RJ, and Chandel NS. We need to talk about the Warburg effect. Nature Metabolism. 2020;2(2):127–9.

27. Warburg O. The metabolism of carcinoma cells. The Journal of Cancer Research. 1925;9(1):148–63.

28. Ma EH, Verway MJ, Johnson RM, Roy DG, Steadman M, Hayes S, et al. Metabolic Profiling Using Stable Isotope Tracing Reveals Distinct Patterns of Glucose Utilization by Physiologically Activated CD8^+^ T Cells. Immunity. 2019;51(5):856–70.e5.

29. Liberti MV, and Locasale JW. The Warburg Effect: How Does it Benefit Cancer Cells? Trends Biochem Sci. 2016;41(3):211–8.

30. Deng Y, Wang ZV, Gordillo R, An Y, Zhang C, Liang Q, et al. An adipo-biliary-uridine axis that regulates energy homeostasis. Science. 2017;355(6330).

31. Puchalska P, and Crawford PA. In: Stover PJ, and Balling R eds. Annual Review of Nutrition, Vol 41, 2021. 2021:49–77.

32. Camarero N, Mascaró C, Mayordomo C, Vilardell F, Haro D, and Marrero PF. Ketogenic HMGCS2 Is a c-Myc target gene expressed in differentiated cells of human colonic epithelium and down-regulated in colon cancer. Mol Cancer Res. 2006;4(9):645–53.

33. Wang YH, Liu CL, Chiu WC, Twu YC, and Liao YJ. HMGCS2 Mediates Ketone Production and Regulates the Proliferation and Metastasis of Hepatocellular Carcinoma. Cancers (Basel). 2019;11(12).

34. Zou K, Hu Y, Li M, Wang H, Zhang Y, Huang L, et al. Potential Role of HMGCS2 in Tumor Angiogenesis in Colorectal Cancer and Its Potential Use as a Diagnostic Marker. Can J Gastroenterol Hepatol. 2019;2019:8348967.

35. Zhang J, Jia PP, Liu QL, Cong MH, Gao Y, Shi HP, et al. Low ketolytic enzyme levels in tumors predict ketogenic diet responses in cancer cell lines in vitro and in vivo. J Lipid Res. 2018;59(4):625–34.

36. Shukla SK, Gebregiworgis T, Purohit V, Chaika NV, Gunda V, Radhakrishnan P, et al. Metabolic reprogramming induced by ketone bodies diminishes pancreatic cancer cachexia. Cancer Metab. 2014;2:18.

37. Sperry J, Condro MC, Guo L, Braas D, Vanderveer-Harris N, Kim KKO, et al. Glioblastoma Utilizes Fatty Acids and Ketone Bodies for Growth Allowing Progression during Ketogenic Diet Therapy. iScience. 2020;23(9):101453.

38. Xia S, Lin R, Jin L, Zhao L, Kang HB, Pan Y, et al. Prevention of Dietary-Fat-Fueled Ketogenesis Attenuates BRAF V600E Tumor Growth. Cell Metab. 2017;25(2):358–73.

39. Lima WG, Martins-Santos ME, and Chaves VE. Uric acid as a modulator of glucose and lipid metabolism. Biochimie. 2015;116:17–23.

40. Yiu A, Van Hemelrijck M, Garmo H, Holmberg L, Malmström H, Lambe M, et al. Circulating uric acid levels and subsequent development of cancer in 493,281 individuals: findings from the AMORIS Study. Oncotarget. 2017;8(26):42332–42.

41. Fini MA, Elias A, Johnson RJ, and Wright RM. Contribution of uric acid to cancer risk, recurrence, and mortality. Clin Transl Med. 2012;1(1):16.

42. Nwosu ZC, Ward MH, Sajjakulnukit P, Poudel P, Ragulan C, Kasperek S, et al. Uridine-derived ribose fuels glucose-restricted pancreatic cancer. Nature. 2023;618(7963):151–8.

43. Skinner OS, Blanco-Fernández J, Goodman RP, Kawakami A, Shen H, Kemény LV, et al. Salvage of ribose from uridine or RNA supports glycolysis in nutrient-limited conditions. Nature Metabolism. 2023;5(5):765–76.

44. Dong Y, Tu R, Liu H, and Qing G. Regulation of cancer cell metabolism: oncogenic MYC in the driver’s seat. Signal Transduction and Targeted Therapy. 2020;5(1):124.

45. Chen C-Y, Chen J, He L, and Stiles BL. PTEN: Tumor Suppressor and Metabolic Regulator. Frontiers in Endocrinology. 2018;9.

46. Zhang X, Zhao H, Li Y, Xia D, Yang L, Ma Y, et al. The role of YAP/TAZ activity in cancer metabolic reprogramming. Molecular Cancer. 2018;17(1):134.

47. Puchalska P, and Crawford PA. Application of Stable Isotope Labels for Metabolomics in Studies in Fatty Liver Disease. Methods Mol Biol. 2019;1996:259–72.

48. Ivanisevic J, Zhu Z-J, Plate L, Tautenhahn R, Chen S, O’Brien PJ, et al. Toward ‘Omic Scale Metabolite Profiling: A Dual Separation–Mass Spectrometry Approach for Coverage of Lipid and Central Carbon Metabolism. Analytical Chemistry. 2013;85(14):6876–84.

49. Koelmel JP, Kroeger NM, Gill EL, Ulmer CZ, Bowden JA, Patterson RE, et al. Expanding Lipidome Coverage Using LC-MS/MS Data-Dependent Acquisition with Automated Exclusion List Generation. Journal of the American Society for Mass Spectrometry. 2017;28(5):908–17.

50. Silva LP, Lorenzi PL, Purwaha P, Yong V, Hawke DH, and Weinstein JN. Measurement of DNA Concentration as a Normalization Strategy for Metabolomic Data from Adherent Cell Lines. Analytical Chemistry. 2013;85(20):9536–42.

51. Puchalska P, Nelson AB, Stagg DB, and Crawford PA. Determination of ketone bodies in biological samples via rapid UPLC-MS/MS. Talanta. 2021;225:122048.

52. Queathem ED, Nelson AB, and Puchalska P. In: Giera M, and Sánchez-López E eds. Clinical Metabolomics: Methods and Protocols. New York, NY: Springer US; 2025:117–31.

53. Baller J, Kono T, Herman A, and Zhang Y. Practice and Experience in Advanced Research Computing 2019: Rise of the Machines (learning). Chicago, IL, USA: Association for Computing Machinery; 2019:Article 96.

54. Bolger AM, Lohse M, and Usadel B. Trimmomatic: a flexible trimmer for Illumina sequence data. Bioinformatics. 2014;30(15):2114–20.

55. Kim D, Langmead B, and Salzberg SL. HISAT: a fast spliced aligner with low memory requirements. Nature Methods. 2015;12(4):357–60.

56. Liao Y, Smyth GK, and Shi W. The Subread aligner: fast, accurate and scalable read mapping by seed-and-vote. Nucleic Acids Research. 2013;41(10):e108-e.

57. McCarthy DJ, Chen Y, and Smyth GK. Differential expression analysis of multifactor RNA-Seq experiments with respect to biological variation. Nucleic Acids Res. 2012;40(10):4288–97.

58. Robinson MD, McCarthy DJ, and Smyth GK. edgeR: a Bioconductor package for differential expression analysis of digital gene expression data. Bioinformatics. 2009;26(1):139–40.

59. The Gene Ontology Consortium. The Gene Ontology Resource: 20 years and still GOing strong. Nucleic Acids Research. 2018;47(D1):D330–D8.

60. Yu G, Wang LG, Han Y, and He QY. clusterProfiler: an R package for comparing biological themes among gene clusters. Omics. 2012;16(5):284–7.

61. Love MI, Huber W, and Anders S. Moderated estimation of fold change and dispersion for RNA-seq data with DESeq2. Genome Biology. 2014;15(12):550.

62. Subramanian A, Tamayo P, Mootha VK, Mukherjee S, Ebert BL, Gillette MA, et al. Gene set enrichment analysis: a knowledge-based approach for interpreting genome-wide expression profiles. Proc Natl Acad Sci U S A. 2005;102(43):15545–50.

63. Ewald J, Zhou G, Lu Y, and Xia J. Using ExpressAnalyst for Comprehensive Gene Expression Analysis in Model and Non-Model Organisms. Current Protocols. 2023;3(11):e922.

64. Chong J, and Xia J. Using MetaboAnalyst 4.0 for Metabolomics Data Analysis, Interpretation, and Integration with Other Omics Data. Methods Mol Biol. 2020;2104:337–60.

